# Advancing Cell Therapies: Single-Cell Profiling, Generation, Expansion, and Gene Delivery in Rhesus Macaque Plasma B Cells

**DOI:** 10.1101/2023.10.29.564645

**Authors:** Rene Yu-Hong Cheng, Shannon Kreuser, Noelle Dahl, Yuchi Honaker, Rupa Soligalla, Christina Lopez, David J. Rawlings, Richard G. James

## Abstract

Engineered long lived plasma cells have the potential to be a new area of cell therapy. A key step in developing this cell therapy is testing in a model with an intact immune system similar to humans. To that end, we have developed methods to purify, expand, and differentiate non-human primate (NHP; *rhesus macaque*) B cells *ex vivo*. We consistently achieved 10-fold expansion of NHP B cells using a readily available commercial supplement. After only seven days in culture, large percentages of cells in NHP B cell cultures were differentiated. These cells expressed surface markers found in human antibody secreting cells (CD38 and CD138) and secreted immunoglobulin G. From single cell transcriptome analysis of NHP, we verified the presence of plasma cell markers commonly shared with humans, and have unearthed less recognized markers such as *CD59* and CD79A. In addition, we identified unique NHP plasma cell markers that are absent in humans including the immune checkpoint molecule *CD274* (PD-L1, Programmed Death-Ligand 1). Furthermore, we found that MHC class I molecules were upregulated in NHP plasma cells, in contrast to the pattern observed in humans. Lastly, we also identified the serotypes (AAVD-J) and established the conditions for efficient transduction of NHP B cells with AAV vectors, achieving an editing rate of approximately 60%. We envision that this work will accelerate proof-of-concept *in vivo* studies using engineered protein-secreting B cells in the NHP model.

## Introduction

Protein or peptide drugs, including monoclonal antibodies (mAbs), are a growing class of therapeutics that now constitute ∼10% of the pharmaceutical market^1^. Although protein drugs have great promise due to their ability to specifically target pathways and cell types that have eluded small molecule inhibitors, they have many drawbacks. These drawbacks can include poor solubility, requirement for mammalian or human post-translational modifications, and relatively short half-life’s *in vivo*. Because of these drawbacks, protein drugs are relatively expensive and can be difficult to manufacture. Recently, we developed a cell-based method to deliver protein drugs, which we have successfully tested in immune-deficient mice. To do this, we generated *ex vivo* differentiated human plasma cells (PCs) and engineered them to produce protein drugs (including bispecific antibodies), and in immuno-deficient mice, we observed long-lasting antibody secretion for a year and potent tumor killing^1–4^.

We, and others have previously shown that human^5^ and murine^6^ PCs can engraft in immune deficient mice (small animals). However, the safety, feasibility and scalability of engineered B cells has not been carefully evaluated in large animal models. Larger animals, such as non-human primates (NHP), more faithfully replicate human immune responses when compared to mice. They provide substantially larger tissues (e.g., bone marrow) for PC residency, more faithful routes of delivery, and the potential for increased longevity for evaluating durability of therapy, all surpassing what is achievable in mice. The rhesus macaque animal model is increasingly used for the preclinical development of HIV vaccines, microbicides and antiretroviral drugs^7^. While conditions for NHP B cell *in vitro* cultures^8^, and suitability for AAV transduction^9^ has been explored, a detailed molecular comparison of these cells with human PCs has not been achieved.

*In vitro* PC products derived from progenitor cells exhibit substantial heterogeneity^1,5^. Examining the genomic profile in NHP cell products is crucial for understanding how likely experiments using these cells are going to mimic a human cell product. Using flow cytometry, Terstappen et al. showed that the canonical human PC marker, CD38 is expressed by NHP bone marrow-derived PCs^10^. Furthermore, single-cell RNA sequencing was used to demonstrate that NHP PCs from blood, bone marrow, and lymph nodes express similar factors as human primary PCs. They identified ICAM2 as a novel biomarker present in primary NHP PCs^11^, which they confirmed in humans. Comparisons between human and NHP can help provide a bridge between preclinical studies in NHPs, and future clinical applications.

In this study, we refined the reagents and methods we previously developed^12^ to establish a comprehensive workflow for expanding and engineering NHP PCs. This included the identification of a range of reagents for isolation, transduction with AAV/lentivirus, expansion, and differentiation of B cells sourced from peripheral rhesus macaque blood mononuclear cells. We anticipate that the level of B cell expansion in this system will enable multiple transfers of PCs at proportions relative to body weight, similar to those employed in previous murine studies of both human and murine PC longevity. Of particular significance, our work addresses a critical knowledge gap by providing molecular insights into the *ex vivo* differentiation of NHP PCs at the single-cell level. We demonstrated that NHP PC cultures exhibit remarkable heterogeneity and identified several potential markers that distinguish different stages of differentiation in NHP PCs. Additionally, through a comparative analysis of NHP single-cell data with previously generated human data, we observed a high degree of conservation in PC genes/markers (such as CD38, SLAMF7, ICAM2, MZB1, and CD59) across primates. However, in opposition to what is observed in human PCs^1,13^, we found that MHC-Class I genes, and CD274 (PD-L1) were significantly increased during differentiation of NHP PCs, which could have implications for evaluating the immunogenicity of engineered NHP PCs. .

## Result

### Workflow for Rhesus Macaque ex vivo PC differentiation and illustrate the heterogeneity of the PC

Human cytokines and human-directed antibodies may not efficiently cross-react with NHP surface proteins. We empirically tested a suite of human reagents (Table S1) with rhesus macaque B cells. The first step in *ex vivo* B cell culture is isolation of B cells from peripheral blood mononuclear cells. We identified an NHP CD20 positive selection kit that enabled enrichment of NHP B cells at ∼80% purity (Flow cytometry panel Table S2, and gating strategy Figure S1, Figure S2A).

Next, we evaluated the efficacy of a human cytokine cocktail that included several T cell dependent cytokines and CpG or a commercial human B cell expansion medium (Figure S2B). Following 7 or 13 days of culture, we found that relative to defined cytokines, the commercial medium elicited greater B cell expansion, and decreased T cell expansion (Figure S2C). To test the effect of seeding density on NHP B cell cultures, we cultured cells for 7 days in the commercial expansion cocktail at high (1.5 million cells mL^-1^) and low (100 thousand cells mL^-1^) densities (Figure 1A-E, S2D-E). We found that NHP B cells grown at low density *ex vivo* exhibited significant increases in total number relative to those grown at high density (Figure 1A; ∼10.5-fold compared to 1.5 fold). We found that B cells grown at low density also exhibited similar viability (Figure S2D) and purity (Figure 1B,C; ∼95% as defined by the CD14-CD3-percentage). Upon evaluating the capacity for differentiation following treatment, we found that the commercial media exhibited fewer CD20^+^ (naive B) cells (Figure 1B,D; <10%), and higher percentages (up to 40%) of CD138^+^ (PCs) cells (Figure 1B,E). Finally, we also detected higher secretion ability per cell with low density culture (Figure S2E), and confirmed that ∼50% of the cells produced IgG via ELISPOT (Top right, Figure 1F). These data indicated that NHP B cells expanded efficiently in commercial cultures and exhibited more rapid differentiation than we previously observed in human B cell cultures (13 days)^1,2^.

**Figure 1:**
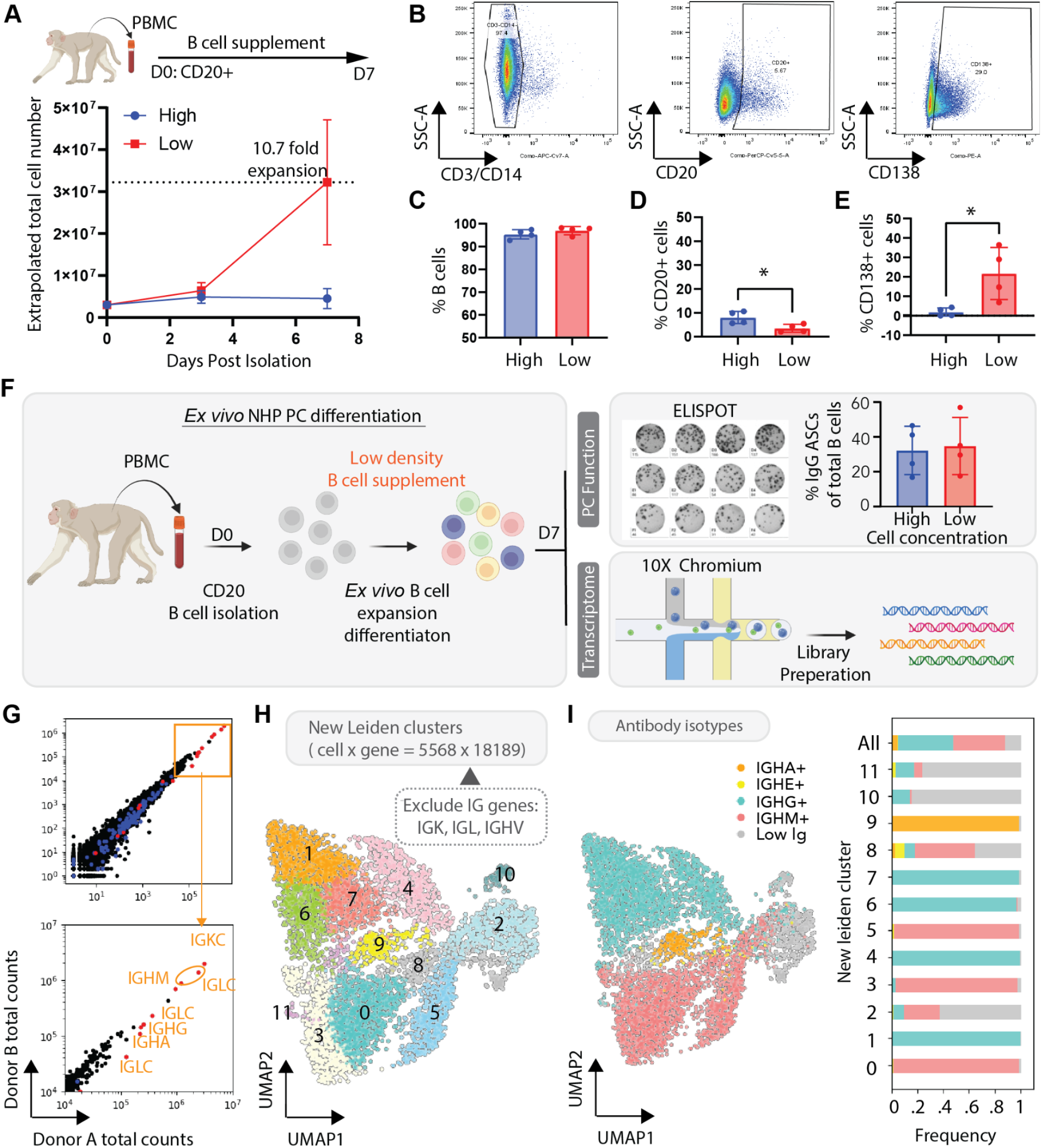
Monkey *ex vivo* differentiated PCs transcriptome with high immunoglobulin expression. **A-E** To assess the impact of cell density, CD20+ NHP B cells were cultured in the commercial B cell medium for 7 days (Plated and maintained 1.5×10^6^ cells/mL versus plated at 2.5×10^5^ cells/mL and maintained at 1×10^5^ cells/mL; 4 donors, n=4). At day 7 following isolation, the cells were analyzed using flow cytometry. **A.** Dynamic of total cell overtime. **B**. Representative flow cytometry plots of CD20, CD38 and CD138 expression. **C.** B cells (CD3-CD14-), **D.** naive B cells (CD20+) and **E.** plasma cells (CD138+). **F.** Schematic workflow of monkey *ex vivo* differentiated PC generation for functional (n=4) and transcriptional profile characterization (n=2). **G.** Scatter plot of RNA total counts from two donors (n=2), red dots represent immunoglobulin constant genes and blue dots represent immunoglobulin variant genes. **H.** UMAP leiden clusters after removing immunoglobulin light chain genes and variant genes. **I.** UMAP projection of antibody isotype. Bar graph of percentage of antibody isotype in each new cluster.

The preliminary phenotype/ELISPOT results suggest efficient *ex vivo* differentiation of NHP B cells into PCs. To more accurately assess heterogeneity in populations of differentiated NHP B cells, we expanded B cells from two donors at low density in commercial media for 7 days. At this point, we fixed the B cells and performed single cell RNA sequencing on the mixed populations using the 10X genomics platform (Figure 1F). Because the NHP genome was not fully annotated, especially within immunoglobulin and the major histocompatibility complex (MHC) genes, we made a workflow to annotate more genes associated with PC differentiation (Figure S3 and gene annotation file in supplementary material).

To characterize NHP *ex vivo* differentiated PCs, we used unsupervised leiden clustering and found the B cell populations to be heterogeneous, with 13 clusters in ∼5600 cells (FigureS4A). We categorized the cells by expression of the highly expressed (Figure 1G) immunoglobulin light chain genes by drawing “gates” at the local minimum between the modes in each distribution (Figure S4B). Using this method, the clustering is primarily driven by the immunoglobulin genes, especially the light chains (IGKC and three IGLC) (Figure S4B). To unearth additional differentiation-related clusters, we excluded the light chains and mapped light chain (LC) independent clusters (Figure 1H). As expected, after reclustering IGKC, IGLCx cells were distributed equally among the resulting 11 clusters (Figure 1H and Figure S4C), which were now distributed into multiple subclusters for each antibody isotype (Figure 1H-I). Upon assessing UMI’s by cluster, we found that clusters 3 and 6 expressed higher mitochondrial genes and lower UMI (Figure S4D). Additionally, clusters 10 and 11 had fewer than 100 cell counts, and few detectable immunoglobulins (Figure 1I) Consequently, we excluded these clusters from further analysis (Clusters 3, 6, 10 and 11). In summary, we predicted that removal of the light chain immunoglobulins and cleanup of non-B cell clusters would enable us to better understand the impact of lower expressed genes, such as transcription factors and surface marker genes.

### Ex vivo differentiated Rhesus Macaque B cells form clusters analogous to that in human B cell cultures

To classify the cell types in the clusters, we looked at genes restricted to specific B cell clusters in human cells^1,13^, including activated B cells (*PAX5, MS4A1, CD79B*, and *MAMU-DPA*), and differentiated B cells (PB/PCs; *IRF4, PRDM1*, and *XBP1*, etc) (Figure 2A-B). Cluster 2 exhibited low immunoglobulin counts and clear expression of activated B cell markers. Additionally, we categorized cluster 8 as a prePB, due to expression of activated B cell markers, and also *IRF4* and *PRDM1*. The remaining clusters exhibited similar expression of human PC markers (Figure 2A-B), with the only obvious markers that qualitatively distinguished PB from PC were cell cycle genes (e.g. *MKI67* and *CDK1*). While some canonical markers *XBP1* and *JCHAIN* (Figure 2B), were uniformly distributed among the NHP PB/PC populations, others like *SLAMF7, CD38,* and *TNFRSF17* (otherwise known as BCMA) were either poorly or differentially expressed within the NHP B cell subsets (Figure 2A-B). After assessment of human PB/PC markers and antibody isotypes, we were able to cluster *ex vivo* differentiated NHP B cells into activated B cells, prePBs, PBs and PCs (Figure 2C).

**Figure 2:**
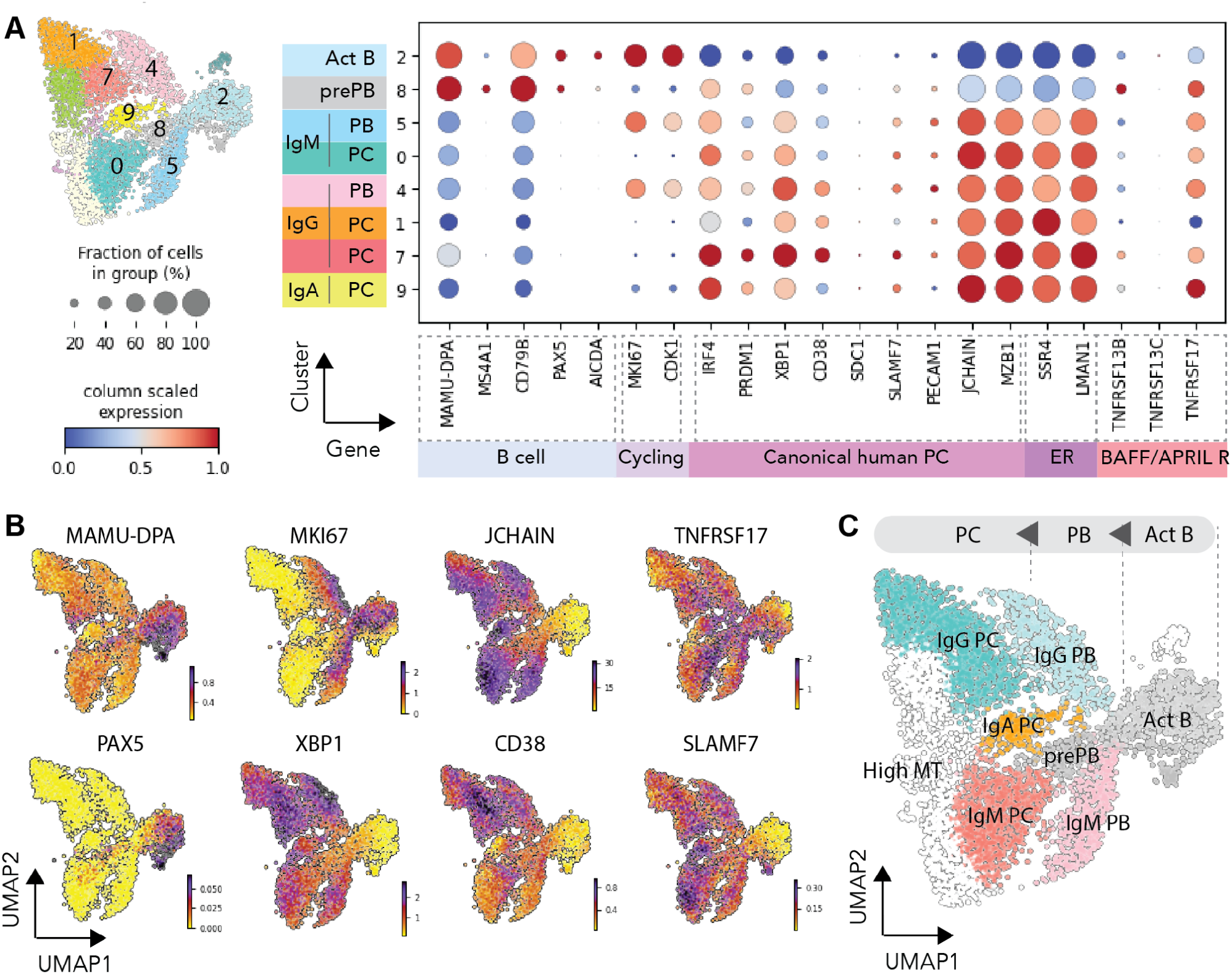
Monkey *ex vivo* differentiated PCs subsets classification and differential expression analysis for PCs. **A.** Dotplot visualization of ex vivo differentiated PCs: subsets are listed on y-axis and genes (features) are listed along the x-axis. Dot size represents percentage of cells in a group expressing each gene; dot color indicates normalized mean expression level in a group. **B.** Heatmap showing expression of representative genes of PC markers (*CD38, JCHAIN*, *XBP1*) and proliferating marker (*MKI67*). **C.** Cluster rename to B cell subsets based on the gene features from figure 2a,b.

### CD59, CD274, CD79A, MAMU-A and MAMU-E were robust Rhesus Macaque PC markers

To identify potential NHP PC markers, we used differential genes expression (DGE) analysis to find genes that are specifically expressed in PCs and/or PBs (Figure 3A). To accomplish this, we performed two comparisons: (1) clusters containing PB/PCs versus more immature CD79B^+^ cells (activated B and prePB), and (2) PCs versus PBs. Similar to previous observations in human cells^1,2,13,14^, we found that *SSR4, MZB1, CD63, JCHAIN, CYBA*, and *TMEM59* were enriched in PBs and PCs relative to CD79B^+^ cells (Figure 3A).^1,2,13,14^ Furthermore, the majority of genes upregulated in PB/PCs were predominantly associated with the endoplasmic reticulum, protein transportation, and protein modification (Figure 3B).

**Figure 3:**
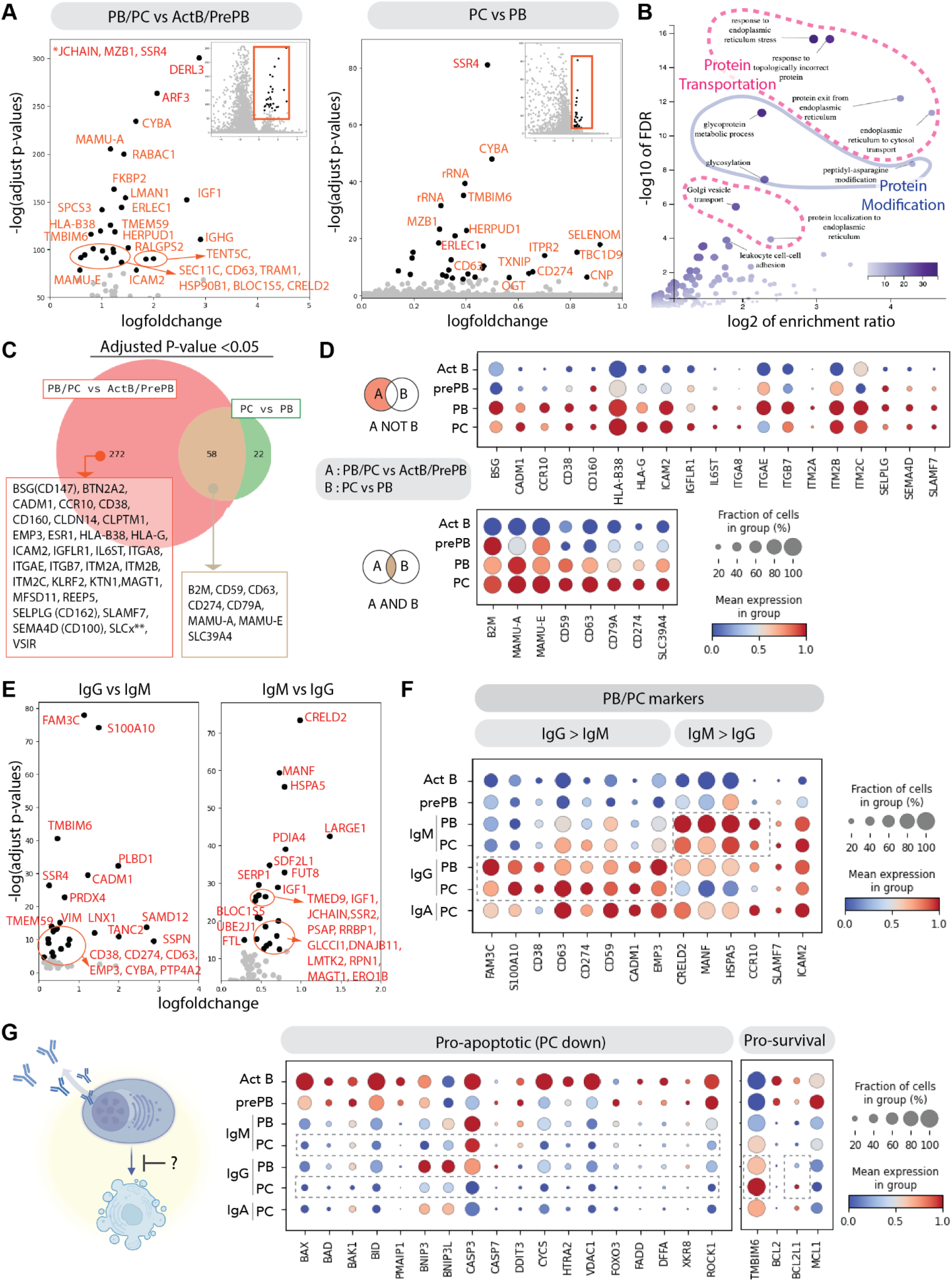
Monkey PCs’ potential markers from DEG. **A.** Volcano plot of differential expression gene, PB/PCs and Activated B/PrePB, PCs vs PBs. Only highlight the top 30 differential genes as black dots. *JCHAIN, MZB1, and SSR4 are infinitely small adjusted p-values, not plotted in the plot. **B.** Volcano plot of enriched gene set from ORA. Heatmap represents the number of genes enriched in each gene set. **C.** Venn diagram of genes upregulated in PB/PCs and PCs, by comparing with ActB/prePB and PBs with adjusted p-value <0.05. Only highlight surface markers or some transmembrane genes. **D.** Dotplot visualization of *ex vivo* differentiated PCs: subsets are listed on y-axis and genes (features) are listed along x-axis. **E.** Volcano plot of differential expression gene, IgG PB/PCs vs IgM PB/PCs (left) and IgM PB/PCs vs IgG PB/PCs (right). Only highlight the top 25 differential genes as black dots. **F.** Dot plot visualization of *ex vivo* differentiated PB/PCs: subsets with different isotypes are listed on y-axis and representative surface marker genes or genes from Fig. 3E are listed along the x-axis. **G.** Schematic cartoon of PC’s anti-apoptotic program. Dot plot visualization of *ex vivo* differentiated PB/PCs: subsets with different isotypes are listed on y-axis and representative anti/pro-apoptotic genes listed along the x-axis.

Because PB/PCs exhibit a global program driven by activation of “master regulator” factors like IRF4, PRDM1 and XBP1^15,16^, we weren’t surprised to find pronounced differences between PB/PCs and CD79B^+^ cells relative to the differences between PC and PBs (Venn diagram showing surface markers, Figure 3C). Several genes enriched in PB/PCs are expressed at similar levels in both cell subsets (Figure 3D, top panel) such as canonical human PC markers, *CD38* and *SLAMF7*, and *ICAM2*, which was previously reported to mark NHP PCs (CD102) (Figure 3D)^11^. In contrast, a subset of surface markers are upregulated in PB/PCs, and are increased in PC relative to PB (Figure 3C, bottom panel e.g., *CD59, CD274, CD79A, MAMU-A*, and *MAMU-E*). In contrast to previous findings in human cells^13^, we observed an upregulation of several MHC-class I molecules in NHP PB/PCs. We previously reported that CD59 is highly expressed in human IgG PCs and correlated with immunoglobulin secretion^2^. Because of its robust expression in NHP PCs, and its correlation with PC function in human cells, we propose that CD59, and possibly other genes in this dataset, are likely to be useful surrogate markers for functional NHP PCs.

To understand whether the identity of the antibody isotype correlated with transcriptional differences, we performed a DGE analysis of IgM and IgG PB/PCs (Figure 3E-F). We found that some canonical PC markers, including *CD38* and some non-canonical markers were expressed at higher levels in IgG, relative to IgM PB/PCs. In contrast, genes associated with endoplasmic reticulum quality control were enriched in IgM (*CRELD2, MANF,* and *HSPA5*) PB/PCs. It is possible that these differentially expressed markers relate to the unique functions of IgM and IgG plasma cells in short- and long-term humoral immunity.

Finally, given the differential expression of apoptotic pathways in long-lived human PCs^13,17,18^, we investigated the presence of apoptotic genes in NHP *ex vivo* differentiated PCs. Similar to the description in human cells, we found that pro-apoptotic genes are generally downregulated during NHP PC differentiation (Figure 3G), with the exception of *BNIP3, BNIP3L*, and *CASP3*, which were upregulated in PBs then downregulated in PCs. Additionally, among pro-survival (anti-apoptotic) genes, we observed upregulation of *TMBIM6* (one of highest IgG DGE genes in FIgure 3E) and *BCL2L1* in IgG PCs (Figure 3G). Collectively, these scRNA sequencing data demonstrate that *ex vivo* derived NHP B cells produced a heterogeneous population that included subsets of PCs with similar transcriptional profiles as long-lived human plasma cells.

### Dynamic expression and kinetic differential analysis of NHP potential PC markers

mRNA splicing can provide supportive information about the progression of cellular differentiation. Upon upregulation of gene expression, unspliced transcripts are often present at a relatively high proportion relative to that observed in steady state conditions^19^. RNA velocity, or the relative ratio of unspliced to spliced RNA can be used to infer upregulation of transcripts in cell subsets within scRNA datasets^19^. As expected, the velocity of IgG, IgM and IgA were increased in PB/PC’s expressing each isotype, although there appeared to be some heterogeneity within each group (top panel, Figure 4A).

**Figure 4.**
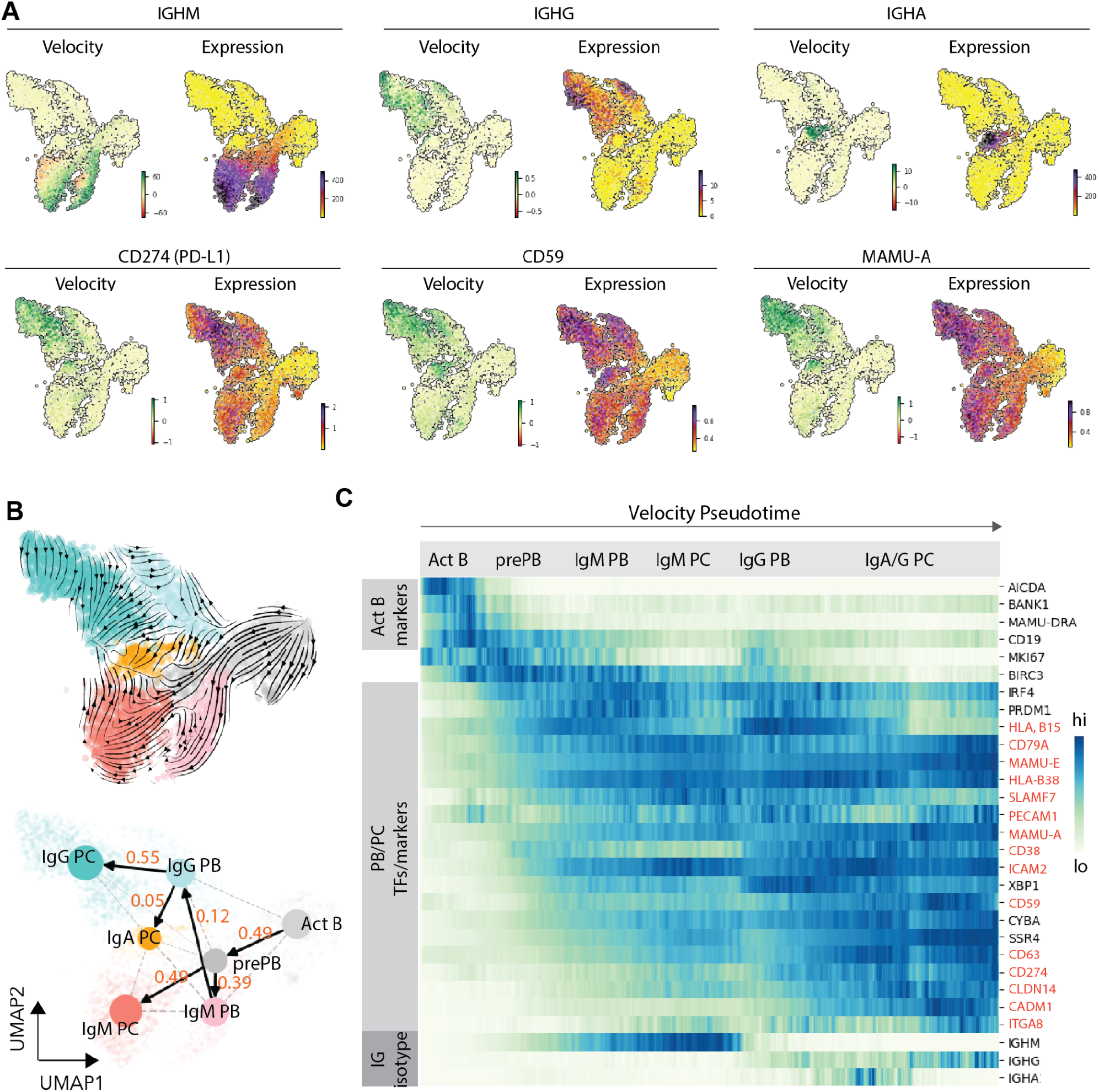
Monkey PC’s dynamic expression. **A.** Heatmap of indicated gene velocity and RNA expression level superimposed on to UMAP. **B.** Velocities derived from the stochastic model and visualized as streamlines in a UMAP-based embedding (Top). PAGA/transition confidence probability from one node to next node (bottom). **C.** Expression heatmap of genes show gene regulation along plasma cell development and genes upregulated in PC from DEG analysis. X-axis along with velocity pseudotime. Y-axis with indicated gene. Potential PC markers in highlighted orange.

In support of the steady-state mRNA data, we found *CD274, CD59* and *MAMU-A* exhibited positive RNA velocity in IgG^+^ PC clusters (Figure 4A). In contrast, genes enriched in activated B cells (e.g. *MAMU-DPA* and *PAX5*), only exhibited positive RNA velocity in non-PC clusters (Figure S5A). Next, we used scVelo, an unsupervised method that leverages RNA velocity to infer the progression of cells along a developmental trajectory, or pseudotime. The pseudotime trajectory generated by the model proceeded from activated cells (right side of UMAP, Figure S5B) to IgG PCs (left side of UMAP, Figure S4B). Using PAGA connection analysis^20^, we calculated the transition probability from one cluster to another (Figure 4B, bottom). The trajectory analysis of activated B cells into antibody-secreting PCs in the NHP system was similar to that observed in human cells^21^. After ordering cells based upon velocity pseudotime, we can see the progression/dynamic of genes of interest along the PC differentiation (Figure 4C, S5C). Similar to the connection analysis, the gene expression trajectories followed canonical B cell differentiation, starting from cells with markers of B cell activation (*AICDA, MHC-II, CD19*, and *BIRC3*), followed by upregulation of transcription factors associated with PB/PCs differentiation (*IRF4*,and *PRDM1*, etc.). Potential PC markers can be classified into three groups based on the timing of the onset: (1) Genes initially expressed in pre-PB (*CD79A, MAMU-E, HLA-B38*, *MAMU-A, CD38,* and *ICAM2*), (2) genes initially expressed in pre-PB/PB (*CD59* and *CD63*), and (3) genes initially expressed in PCs (*CD274, CLDN14, ITGA8,* and *CADM1*). By applying kinetic differential analysis (Figure S5D), *CD274*, was also identified as being differentially upregulated in IgG PCs, validating its importance as a potential marker for NHP PCs.

### Primate: Rhesus Macaque PCs in comparison with human PCs

Previously we used CITE-seq to characterize the *ex vivo* differentiated Human PCs with CD38, CD138, IgM oligo-conjugated antibodies^1^. We used these markers to categorize human B cells into subsets based on surface expression: Activated B (IgM^hi^CD38^-CD138-^), pre-PB (IgM^hi^CD38^lo^CD138^-^), PB (IgM^lo^CD38^+^CD138^-^) and PC (IgM^lo^CD38^+^CD138^+^) (Figure 5A, left UMAP), which we validated via quantification of the B cell and PC TFs, *PAX5* and *XBP1,* respectively (Figure 5B). In PCs, the majority of antibody isotype is IgG (Figure 5A, right UMAP). *Ex vivo* differentiated NHP PCs primarily expressed IgG, but exhibited higher proportions of IgM than we observed in human cultures. Additionally, the NHP PCs exhibited similar maturity (e.g., PC markers in Figure 3, ER/ Golgi features in Figure 3B, ELISPOT in Figure 1F) following 7 days of culture as humans achieved in 13 days.

**Figure 5.**
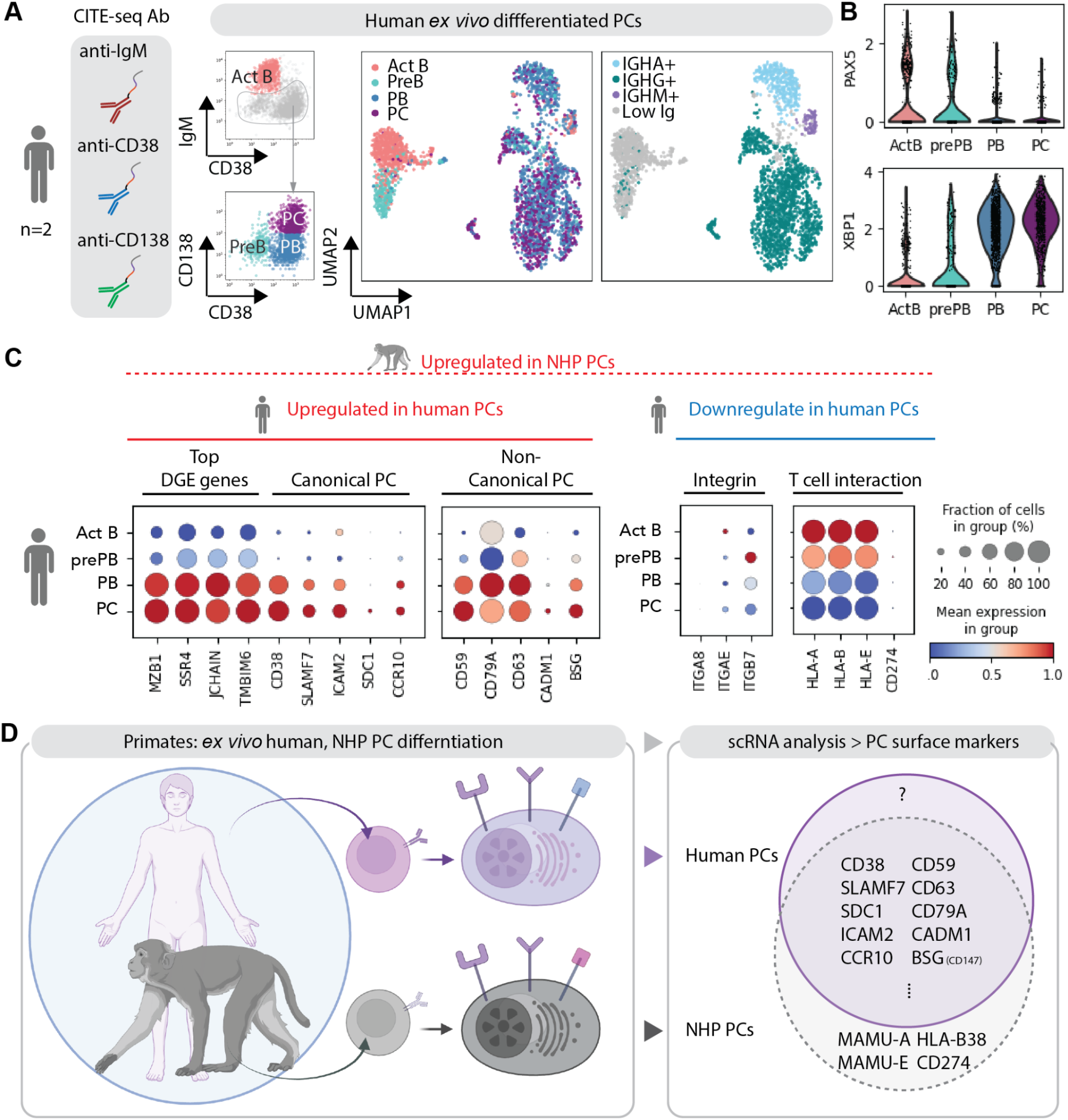
Comparison of human and monkey PC makers by scRNAseq analysis. **A.** *Ex vivo* differentiated human PCs with CITESEQ. Classification of B cell subsets categorized by the indicated protein markers: IgMhiCD38lo (ActB), IgMloCD38lo (PrePB), CD38hiCD138lo (PB), and CD38hiCD138hi (PC). B cell subsets and isotype superimposed on to UMAP projection, day 13 B cells (*n* = 2897) from two biological replicates. **B.** Violin plot of indicated genes. **C.** Dotplot visualization of *ex vivo* differentiated human PC: subsets are listed on y-axis and representative genes from Fig. 3D are listed along the x-axis. **D.** Schematic cartoon summary of human PC markers comparing with monkey PC markers from scRNA analysis

We proceeded to compare the NHP PC markers we identified (Figure 3) with those in human PCs. As anticipated, canonical markers such as *CD38* and *SLAMF7* exhibited higher expression in PB/PC stages. In both species, we found that *CD38* exhibited higher expression in IgG PCs (Figure 5C, S6) and the paramount PC marker, CD138 (*SDC1*) was expressed at higher levels in PCs, but exhibited low RNA counts in all subsets (Figure 2A, 5C and^1^). *CCR10*, canonically expressed in mucosal IgA PCs, was detected specifically in IgA PCs in both species (Figure 5C, S6). Finally, we found several understudied markers, including *CD59*, *CD79A, CD63, CADM1 and BSG* (CD147), to be upregulated in both human and NHP PCs, suggesting further research is justified to clarify potential roles for these genes in PC biology. Of particular interest was the behavior of MHC-class I genes. *HLA-A, HLA-B* and *HLA-E*, which while highly expressed in all human B cell subsets, are downregulated in PCs (Figure 5C). This results directly contrasts our observations in NHP PCs (Figure 3G), whereby the MHC-class I genes are upregulated during PC maturation. Additionally, *CD274* (PD-L1), which is dramatically upregulated in NHP PCs, was hardly detectable in human PCs (Figure 5C, left panel). Collectively, these data imply that while the transcriptional signatures of *ex vivo* derived PCs are well conserved between NHP and humans, there may be substantive differences in expression of T cell regulators between the species.

### Efficient transduction of NHP B cells with the AAV serotypes

To assess the potential for transduction of NHP B cells with recombinant AAV vectors, we isolated B cells, cultured briefly and exposed the cells to a panel of self-complementary AAV vectors. We assessed a series of AAV with alternative serotypes, each carrying a GFP expression cassette to permit efficient tracking of transduced cell populations. Following exposure to AAV, B cells were maintained in culture for nine more days prior to assessment of GFP positivity in CD3-CD14-cells. In a volume matched experiment (20% AAV by volume), we found that AAV D-J serotypes exhibited the most efficient transduction (Figure S7A-B, Table S3). To rule out the impact of titer, we directly compared transduction with the 2.5 (control) and D-J serotypes using the same titers. We found that 2.5 and D-J exhibited similar transduction efficiencies at day2, but D-J increased the efficiencies to 60% at day 4 and remained high percentages at late time points (Day 10; Figure 6B). Next, to determine whether AAV D-J is useful for delivery of a candidate, physiologically relevant, payload in NHP B cells, we transduced NHP B cells with single-stranded AAV expressing the cytokine BAFF cis linked to GFP. While the transduction rates using the BAFF AAV vector were slightly lower than the self-complementary GFP vector (Figure 6C), we detected high levels of BAFF secretion (Figure 6D) in NHP B cells. We also tested whether transduction was different in B cells cultured at different densities. We found that NHP B cultured at low density were transduced at higher rates than those cultured at high densities (Figure 6E), and produced more BAFF in culture (Figure 6D). In conclusion, because AAV DJ can reproducibly be produced at higher titers, we predict that the DJ serotype will be more useful than 2.5 for transduction of NHP B cells.

**Figure 6.**
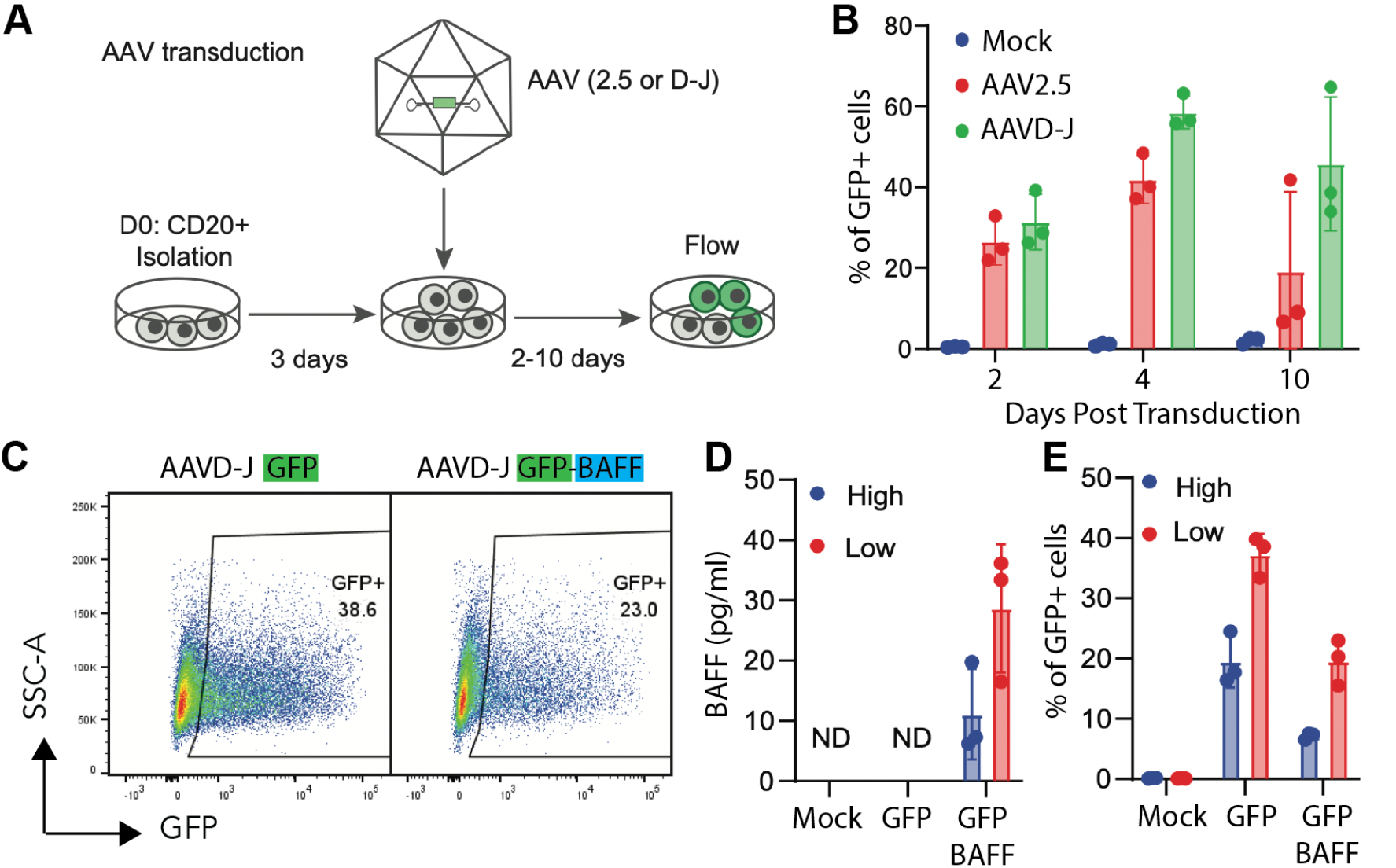
Efficient transduction of NHP B cells with the AAV. **A.** Diagram of the AAV and transduction and culturing protocol. **B**. NHP CD20+ B cells were cultured for 3 days before transduction with the indicated AAV pseudotypes expressing GFP (MOI = 4×104). Transduction efficiency was quantified by flow cytometry at the indicated time points. **C-E** NHP CD20+ B cells grown at high (1.5×106 cells/mL) or low density (2.5×105 cel/m) were transduced with either AAVD-J GFP or AAVD-J GFP-BAFF (3 donors, n=3). **C.** Representative flow plot of GFP and GFP-BAFF transduced cells. **D.** Bar plot of BAFF secretion, was quantified four days post-transduction. **E.** Bar plot of percentage of GFP+ cells with indicated cell density high or low and AAV.

### Assessing Lentiviral transduction of NHP B cells for possible use for gene delivery

B cells from multiple species, as well as rhesus macaque cells from several hematopoietic lineages, have been historically difficult to transduce with gamma-retroviral or lentiviral vectors^22^. Using the XHIV/SHIV packaging system^23^, we made both VSVg and Cocal pseudotyped lentiviral vectors with a GFP payload. After 2 days in culture, each vector was added to NHP B cells (MOI 40). Following vector delivery, B cells were cultured for 11 days in the commercial media, and the transduction was quantified using flow cytometry (Figure S8A). At no point did we observe greater than 2% of the population expressing GFP with either vector (Figure S8B). We also tested lentivirus pseudotyped with a modified version of the Nipah F and G proteins that specifically binds to the CD20^24^ receptor, which is highly expressed on NHP B cells. When using this modified Nipah pseudotype, we found that the percentage GFP positive cells was also less than 3% (Figure S8C), although the mean fluorescent intensity of the GFP positive population was substantially greater than that seen with the VSVg and Cocal pseudotypes(Figure S8C). These results demonstrate that lentiviral transduction of NHP B cells remains a hurdle. However, our findings suggest that using a receptor-targeted Nipah pseudotype that is more specific to NHP CD20 or to a different receptor highly expressed on NHP B cells ultimately could facilitate efficient transduction of NHP B cells.

## Discussion

An important proof of concept experiment for developing a future human plasma cell-based therapy is assessment of plasma cell survival and production capacity in an immunocompetent system. Although NHP models have been used extensively to study the response to vaccines^25^ and PC longevity^26^, we are unaware of comprehensive studies optimizing expansion/differentiation of NHP PCs, describing their heterogeneity, or comparing them to human PCs. To address this challenge, we developed methods to purify, expand, and differentiate *rhesus macaque* B cells *ex vivo.* We achieved a 10-fold expansion of NHP B cells and observed efficient differentiation into antibody secretion cells (∼90 % immunoglobulin high cells). Using single cell RNA sequencing, we confirmed that after seven days in culture, the majority of differentiated NHP B cells express markers associated with antibody secreting cells, including transcription factors, genes regulating protein secretion and immunoglobulin molecules. Furthermore, we identified several surrogate NHP PC markers (CD59, CD274) and compared the transcriptional profiles of NHP and human *ex vivo* derived PCs. Finally, we identified the serotypes (D-J) and conditions necessary for efficient transduction of NHP B cells with AAV vectors, which we hope will be used in homology-directed repair studies to generate cells producing biologic drugs.

Of particular significance, our observations revealed an upregulation of several MHC-class I molecules in NHP (*B2M, HLA-A, HLA-E, HLA-G*) PB/PCs. While it is well-established that MHC-class II molecules are downregulated in human PCs, the expression of MHC-class I genes by PCs in other species has received relatively limited characterization^27–29^. From our human PC single cell RNAseq data (Figure 5) and that generated by other groups^13^, MHC-class I genes are downregulated in human PCs. We also observed upregulation of PD-L1 (CD274) in NHP, but not human PCs^1^. PD-L1, a cell surface protein expressed by various cell types, including cancer cells. PD-L1 expression by cancer inhibits cytotoxic T cell function and promotes immune evasion^30–32^. Because MHC-I and PD-L1 are likely to drive opposing effects on T cell function, it will be important to understand whether these differences between NHP and human PCs have functional impacts on the presentation of foreign antigens by NHP PCs, and the impact on T cell surveillance. It should be noted that PCs express antibody variable chains, which contain antigens that are not thymically educated and would be perceived as foreign. Because NHP can produce long-lived PCs^26^ while expressing antibody variable chains, it is unlikely that expression of MHC-I by NHP PCs leads to PC removal by the immune system. However, this is an area that warrants additional evaluation.

To address the question of how many PCs will be required in primates to generate a neutralizing dose of antibody or relevant levels of alternative therapeutic proteins, there are several variables to be considered: half-life of the protein, effective dose, the production of protein per plasma cell and the engraftment capacity of the plasma cell product. Fortunately, for most clinical mAbs, we know the half-life and the required neutralizing dose. For example, the recently developed half-life extended version of palivizumab, MEDI8897, and standard palivizumab have half-lives of approximately 100 days (2400 hours^33^) and 20 days respectively^34^. Given the half-life (t_1/2_) and volume of displacement (V_d_; 880 mL in a 15 kg adult primate subject), one can estimate the rate of drug clearance (CL). Likewise, using the constant rate infusion equation, given a desired steady-state concentration (*C_ss_*), we can calculate the required daily production. The trough concentration required for neutralization of pathogens is known for many mAbs including CR6261 in influenza (∼10 mg/mL^35^), VRC01 in HIV (∼10 mg/mL^36^) and MEDI8897 in RSV (∼8 mg/mL^33^). Applying these equations to MEDI8897, the approximate daily production (*R_0_*) required for RSV neutralization is approximately 56 mg/day. Because fully differentiated long-lived plasma cells are estimated to produce between 50 pg/cell/day^37^, we predict that stable engraftment of 5 million primate PCs would be sufficient to produce a prophylactic dose of antibody at steady-state.

Upon successful development of engineered primate PCs, we expect that production of a non-clinical cell therapy product could be multiplexed and delivered singly or repeatedly into an autologous donor recipients. Using leukapheresis, approximately 5 billion nucleated cells can be acquired from a single donor in a single extraction^38^ and ∼10% of these cells are expected to be B cells. Based on the rates of expansion observed here (10-fold), following expansion and differentiation, we predict that it will be possible to generate ∼5 billion engraftable plasma cells from a single primate leukapheresis extraction. Given the number of cells we typically transplant into mice to elicit therapeutically relevant and stable antibody secretion^5^, approximately 2.5 billion cells will be required to generate similar antibody levels in an autologous primate transfer. In summary, these results suggest that proof-of-concept experiments using one or more doses of engineered primate plasma cells are likely to be feasible with minor adjustments to the protocol presented here.

## Method

### B cell isolation

Frozen rhesus macaque (*macaca mulatta*) PBMCs were obtained from a commercial source (BioIVT). Using an NHP specific CD20 positive selection isolation kit (Miltenyi Biotec; Table S1), we isolated CD20+ cells.

### B cell culture conditions

Unless otherwise stated, all cells were cultured in a base media composed of Iscove’s modified Dulbecco’s medium (IMDM) (Gibco) supplemented with 10% FBS (Omega Scientific, FB-11), 1% Glutamax (Gibco), and 55 mM beta-mercaptoethanol (BMe). Where noted cells were also cultured in ImmunoCult-XF (ICXF) base media (StemCell, 10981) The commercial expansion cocktail we used was ImmunoCult™-ACF Human B Cell Expansion Supplement (StemCell,10974). For the defined B cell expansion and differentiation cocktails, the compositions are as follows. Phase I: 100 ng/mL recombinant human MEGACD40L (Enzo Life Sciences), 1 mg/mL CpG oligodeoxynucleotide 2006 (Invitrogen), 250 ng/mL IL-2 (PeproTech), 50 ng/mL IL-10 (PeproTech), and 10 ng/mL IL-15 (PeproTech), and 50ng/mL IL-21 (PeproTech, 200-21). Phase II: IL-2 (250 ng/mL), IL-6 (50 ng/mL, PeproTech), IL-10 (50 ng/mL), and IL-15 (10 ng/mL). Phase III: IL-6 (50 ng/mL), IL-15 (10 ng/mL), and human interferon-a 2B (15 ng/mL, Sigma-Aldrich). All cell counts were done manually using a hemocytometer and trypan blue exclusion.

### Flow cytometry

Flow cytometry was run on an LSR II flow cytometer (BD Biosciences) and data were analyzed using FlowJo software (Tree Star). All cells were fixed using Fix/Perm (BD Biosciences) for 20 min prior to intracellular staining and/or analysis. The flow cytometry antibody panels (Table S2) and gating schemes (Figures S1) are detailed in supplemental figures.

### ELISpot and ELISAS

Membranes were coated with an anti-human IgG capture antibody (Thermo Fisher Scientific, A18813). Cells were then added to the plate in duplicate in 2 fold serial dilutions and incubated for 24hrs prior to detection using an alkaline phosphatase-linked anti-human IgG detection antibody (Jackson, 109-055-008). Protein secretion levels in culture supernatant were quantified via ELISA for human IgG (Invitrogen, 50-112-8849) and human BAFF (R&D Systems, DY124-05) using protocols recommended by the manufacturer.

### Virus production and transduction

All viruses and all pseudotypes, AAV and lentivirus, were made in house previously published methods (see Supplementary note for description of the CD20-pseudotyped Nipah vector). GFP-BAFF was previously published in King et al. AAV transduction was done on day three of culture. First cells were replated at 1.5×10^6^ cells/mL (unless otherwise noted) in IMDM + Bme + Glutamax + commercial supplement without FBS, then virus was added to culture at 20% by volume unless indicated otherwise and incubated at 37°C. After two hours FBS was added back to the media for a final concentration of 10% and returned to 37°C. The next day the media volume was doubled in order to minimize any cell death caused by the addition of the AAV. For Lentivirus transduction, cells were replated at 1.5×10^6^ cells/mL IMDM alone, and virus was added. Cells were spinoculated for 30min at RT at 400xg, then returned to 37°C for six hours. After six hours cells were spun down for 5 min at 400xg, half the media volume was taken, and IMDM was added back with two times the concentration of FBS, BMe, Glutamax, and commercial supplement and returned to 37°C.

### Single Cell RNA Sequencing

NHP Single cell cDNA libraries were created using the 10x genomics platform (3’ chemistry V3.1 dual index kit, 10x genomics). Human cells were labeled with oligo-conjugated antibodies for tracking surface expression and sample identity labeling using the Biolegend Totalseq-B protocol. We prepared libraries using the 10X genomics platform (Single-Cell v3.1 with feature barcode dual index kit, 10x genomics). Both NHP and human libraries from cell surface tags and transcripts were evaluated by tapestation (Agilent) before sequencing. Finally, libraries were sequenced with NextSeq 1000/2000 kit (Illumina) .

### Single-cell RNA-seq analysis

Fastq files were processed by CellRanger and Velocyto based on the customized *Macaca mulatta* gene annotation (Figure S2) . The h5 file is then further analyzed by a python script. And following analysis including normalized, trajectory inference (data clustered using the Leiden-graph-based algorithm and reconstructed the lineage relations by PAGA), and other hierarchical clusterings, dimensional reduction, and cell clustering is analyzed by python package scanpy. Cell velocity and velocity pseudotime is analyzed by python package scVelo.

### Data availability

The single cell RNA sequencing data are available at GSE247023.

## Supplemental data

Table S1: Reagents and antibodies

Table S2: Staining panel

Table S3: AAV transduction with different titer

Figure S1-S8

Supplemental note: Production of anti-CD20 pseudotyped lentivirus

---

**Table S1:** Reagents and antibodies.

| Reagent | Company | Cat |
| --- | --- | --- |
| NHP CD20 Isolation Kit | Miltenyi | 130-091-105 |
| IMDM | Gibco | 12-440-053 |
| ICXF | StemCell | 10981 |
| StemCell B-cell supplement ACF | StemCell | 10974 |
| Multimeric CD40L | Adaptogen | AG-40B-0010 |
| CpG oligodeoxynucleotide 2006 | Invitrogen | tlrl-2006-5 |
| IL-2 | PeproTech | AF-200-02 |
| IL-10 | PeproTech | 200-10 |
| IL-15 | Miltenyi | 130-095-765 |
| IL-21 | PeproTech | 200-21 |
| IL-6 | PeproTech | 200-06 |
| IFN alpha 2B | Proteintech | HZ-1072 |
| AF350 amine reactive dye | Invitrogen | A10168 |
| anti-CD3 APC-Cy7 clone:SP34-2 | BD Bioscience | 557757 |
| anti-CD3 A700 clone:SP34-2 | BD Bioscience | 557917 |
| anti-CD4 BV605 clone:L200 | BD Bioscience | 562843 |
| anti-CD14 APC.Cy7 clone:MoP9 | BD Bioscience | 557831 |
| anti-CD20 PerCPCy5.5 clone:L27 | BD Bioscience | 340955 |
| anti-CD31 BV605 clone:WM59 | Biolegend | 303122 |
| anti-CD31 PE clone:WM59 | BD Bioscience | 555446 |
| anti-CD38 mFluor450 clone:OKT10 | Caprico<br>Biotechnologies | 1008144 |
| anti-CD56 PE clone:MY31 | BD Bioscience | 347747 |
| anti-CD138 PE clone:DL-101 | Biolegend | 352306 |
| anti-IgG AF700 clone:G18-145 | BD Bioscience | 561296 |
| anti-IgM FITC clone:G20-127 | BD Bioscience | 555782 |
| IgG capture antibody (ELISpot) | Invitrogen | A18813 |
| IgG AP detection antibody (ELISpot) | Jackson | 109-055-008 |
| IgG (Total) ELISA | Invitrogen | 885055088 |
| BAFF DuoSet ELISA | R&D Systems | DY124-05 |

**Table S2:**
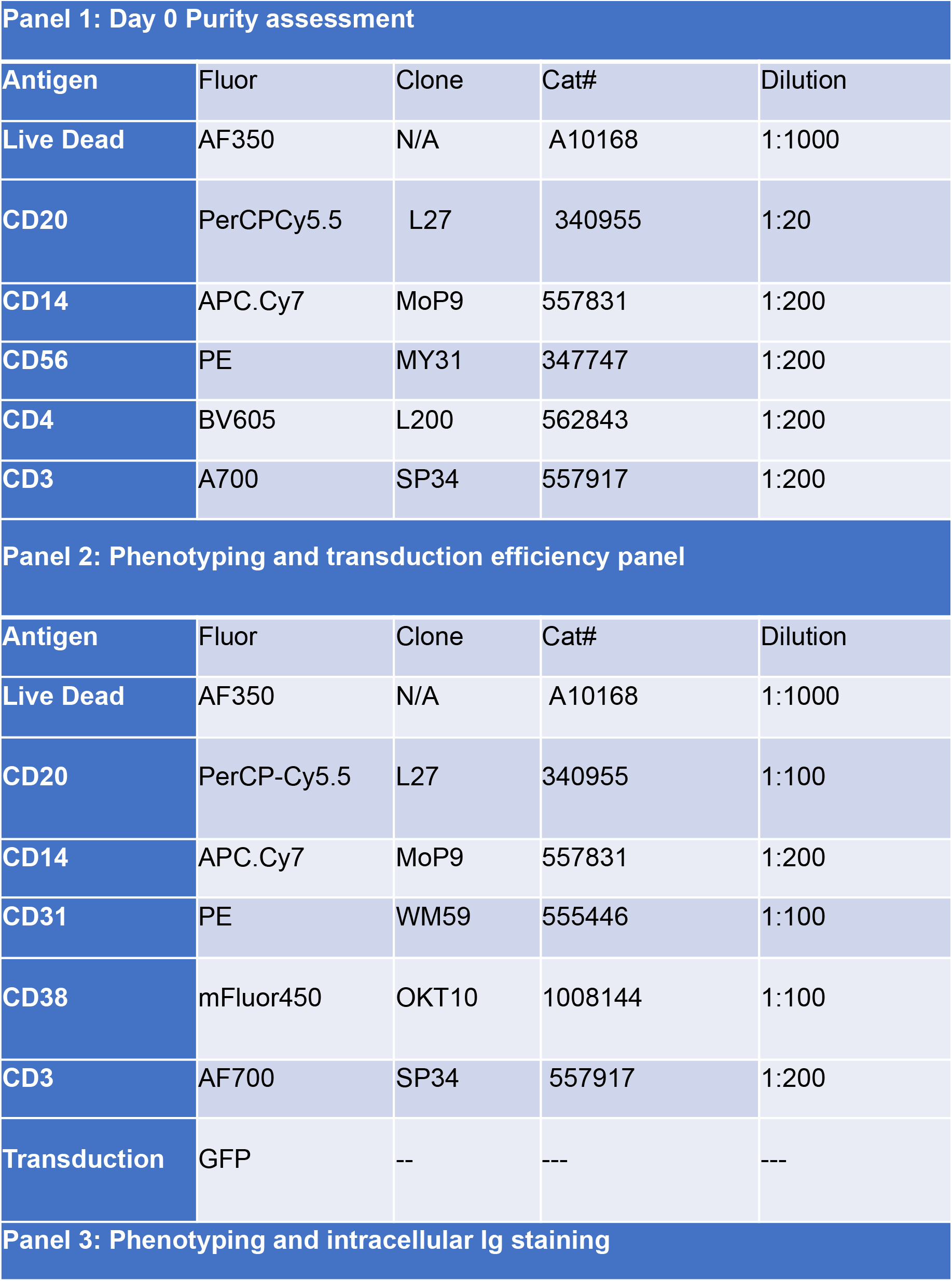
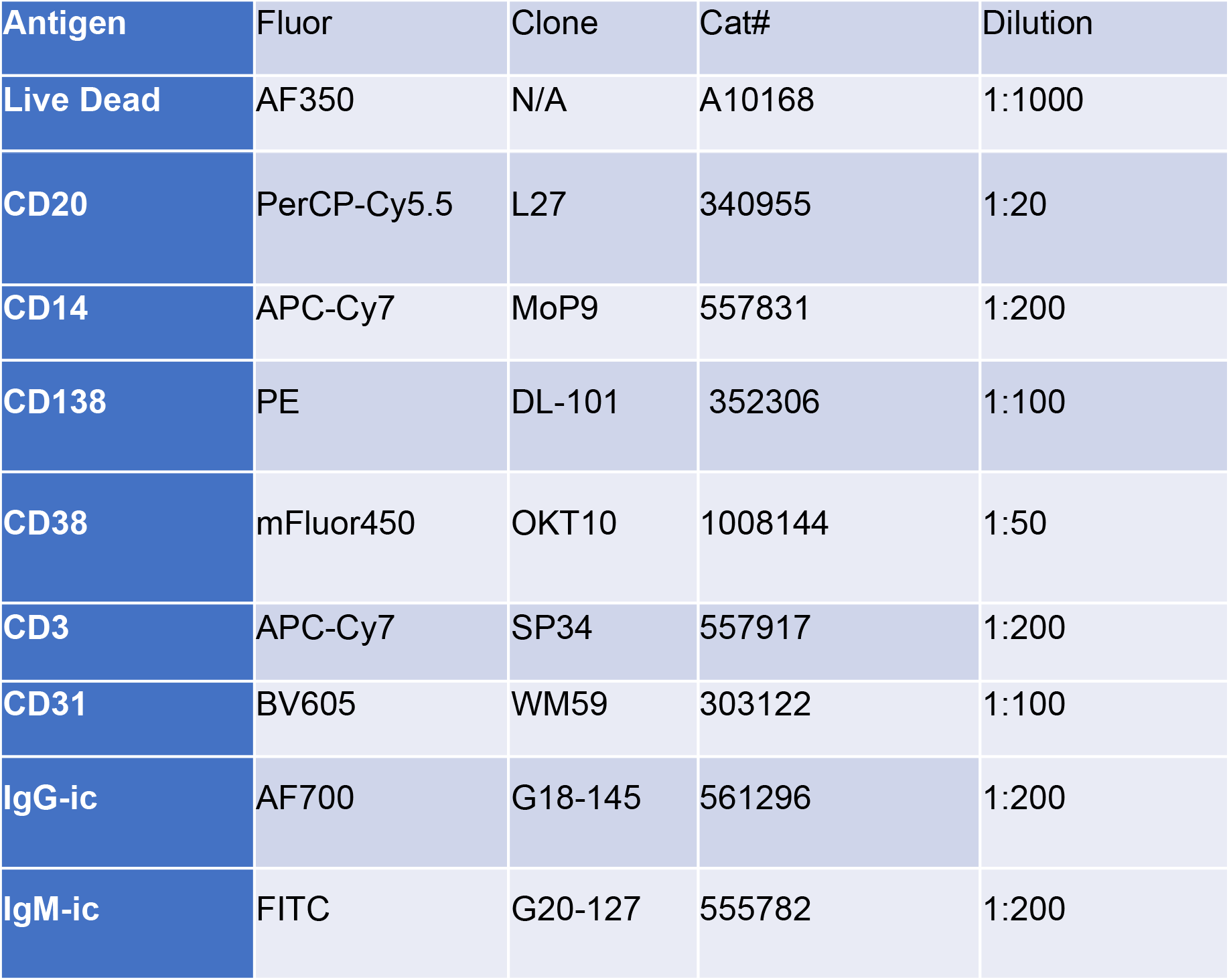
Staining Panel.

| Panel 1: Day 0 Purity assessment |  |  |  |  |
| --- | --- | --- | --- | --- |
| Antigen | Fluor | Clone | Cat# | Dilution |
| Live Dead | AF350 | N/A | A10168 | 1:1000 |
| CD20 | PerCPCy5.5 | L27 | 340955 | 1:20 |
| CD14 | APC.Cy7 | MoP9 | 557831 | 1:200 |
| CD56 | PE | MY31 | 347747 | 1:200 |
| CD4 | BV605 | L200 | 562843 | 1:200 |
| CD3 | A700 | SP34 | 557917 | 1:200 |
| Panel 2: Phenotyping and transduction efficiency panel |  |  |  |  |
| Antigen | Fluor | Clone | Cat# | Dilution |
| Live Dead | AF350 | N/A | A10168 | 1:1000 |
| CD20 | PerCP-Cy5.5 | L27 | 340955 | 1:100 |
| CD14 | APC.Cy7 | MoP9 | 557831 | 1:200 |
| CD31 | PE | WM59 | 555446 | 1:100 |
| CD38 | mFluor450 | OKT10 | 1008144 | 1:100 |
| CD3 | AF700 | SP34 | 557917 | 1:200 |
| Transduction | GFP | -- | --- | --- |
| Panel 3: Phenotyping and intracellular Ig staining |  |  |  |  |

| Antigen | Fluor | Clone | Cat# | Dilution |
| --- | --- | --- | --- | --- |
| Live Dead | AF350 | N/A | A10168 | 1:1000 |
| CD20 | PerCP-Cy5.5 | L27 | 340955 | 1:20 |
| CD14 | APC-Cy7 | MoP9 | 557831 | 1:200 |
| CD138 | PE | DL-101 | 352306 | 1:100 |
| CD38 | mFluor450 | OKT10 | 1008144 | 1:50 |
| CD3 | APC-Cy7 | SP34 | 557917 | 1:200 |
| CD31 | BV605 | WM59 | 303122 | 1:100 |
| IgG-ic | AF700 | G18-145 | 561296 | 1:200 |
| IgM-ic | FITC | G20-127 | 555782 | 1:200 |

**Table S3:**
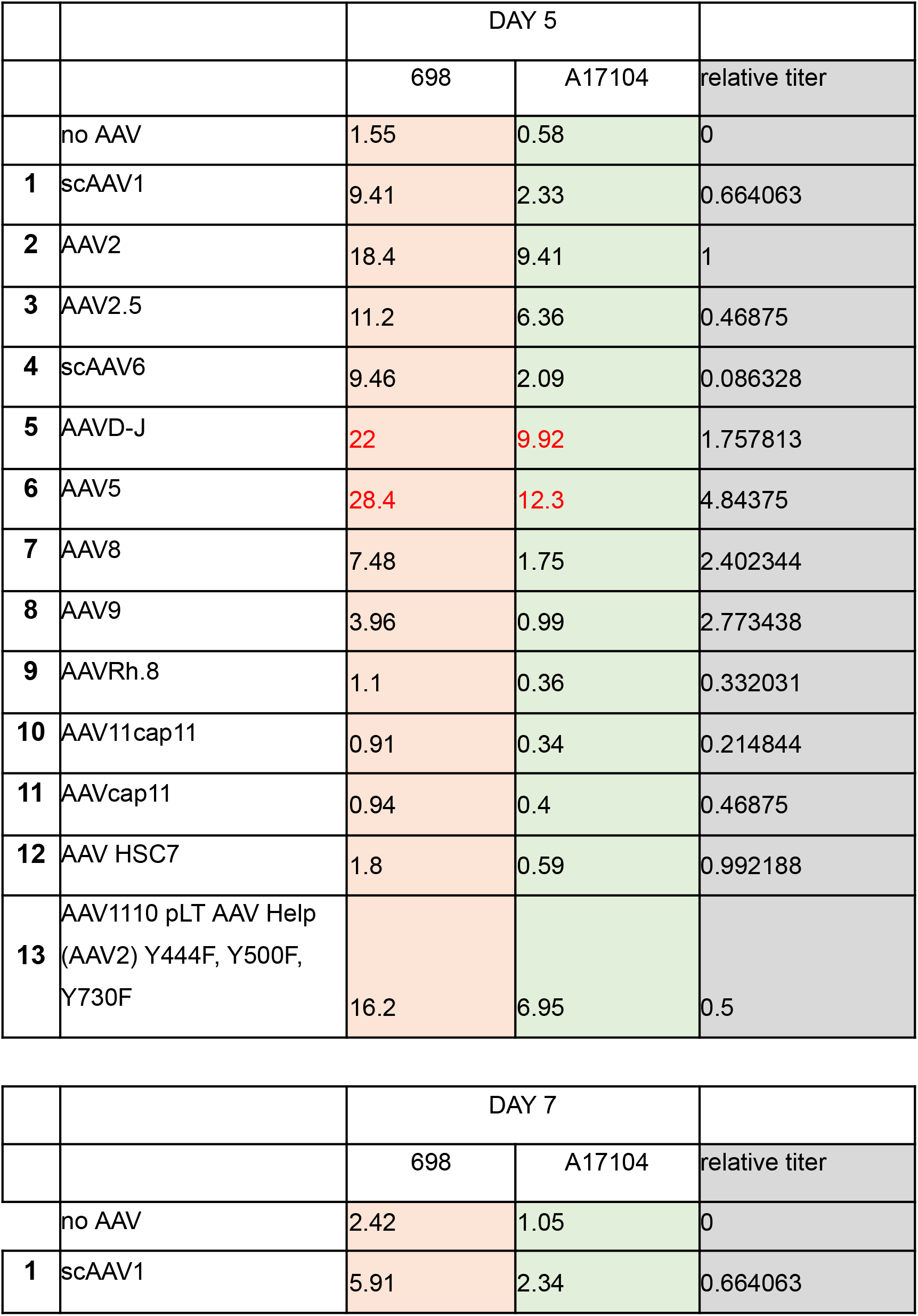
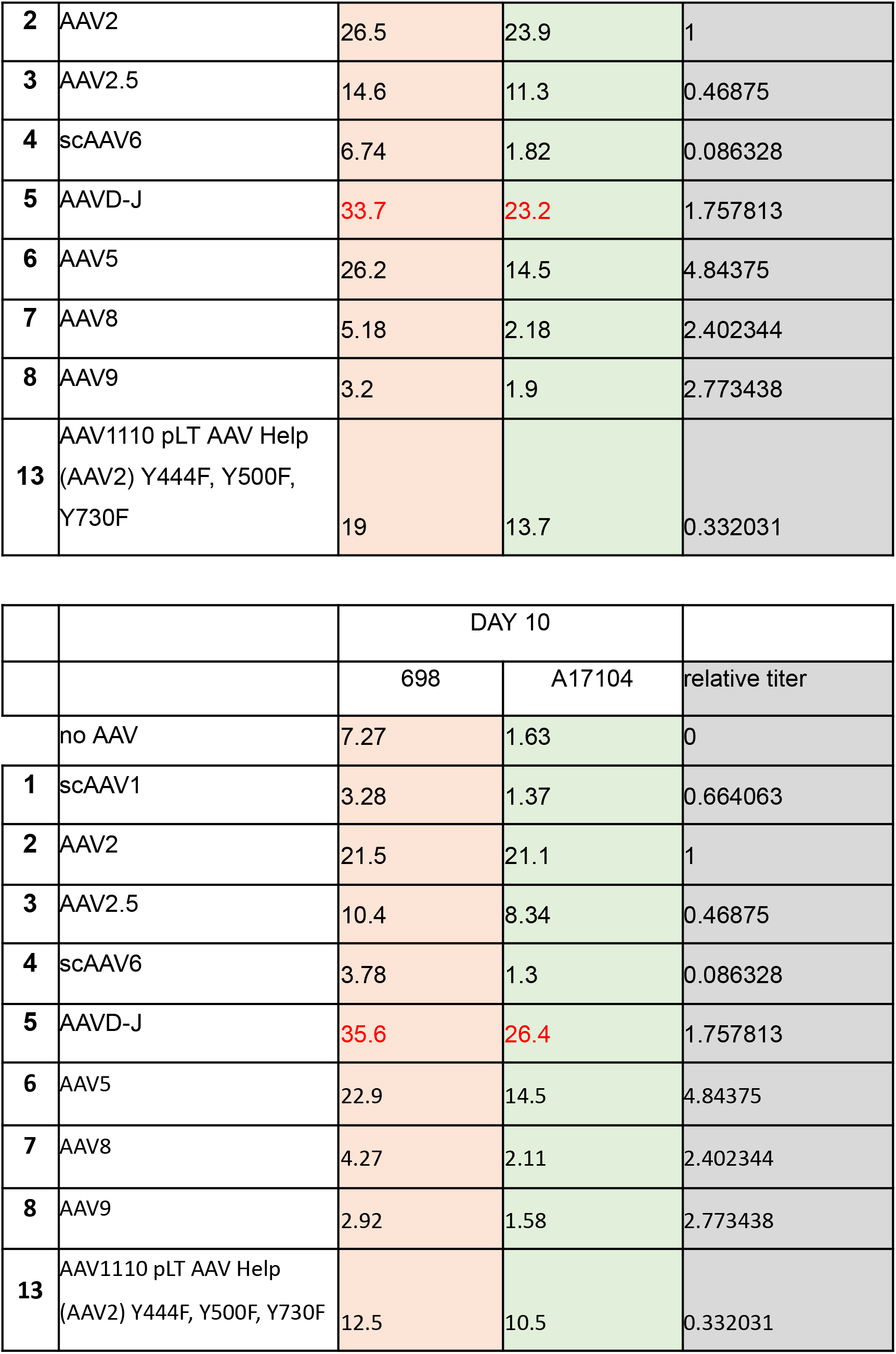
AAV transduction with different titer.

**Figure S1:**
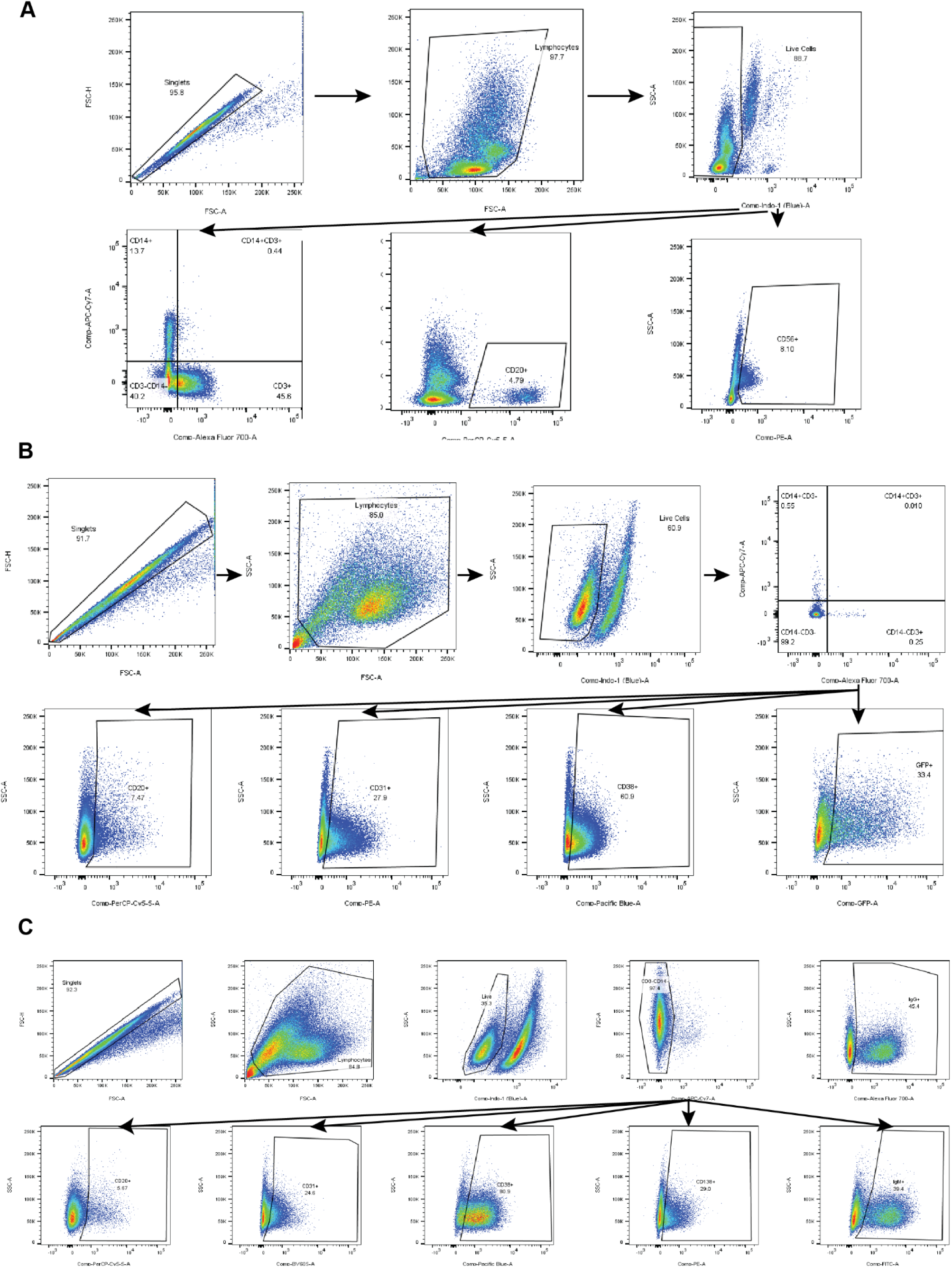
Flow cytometry gating strategy. **A.** Gating scheme for flow cytometry Panel 1 (Table S2) to determine purity at the time of isolation. **B.** Gating scheme for flow cytometry Panel 2 (Table S2) to determine B cell phenotype and transduction efficiency through GFP expression. **C.** Gating scheme for flow cytometry Panel 3 (Table S2) to determine B cell phenotype and intracellular Ig production.

**Figure S2:**
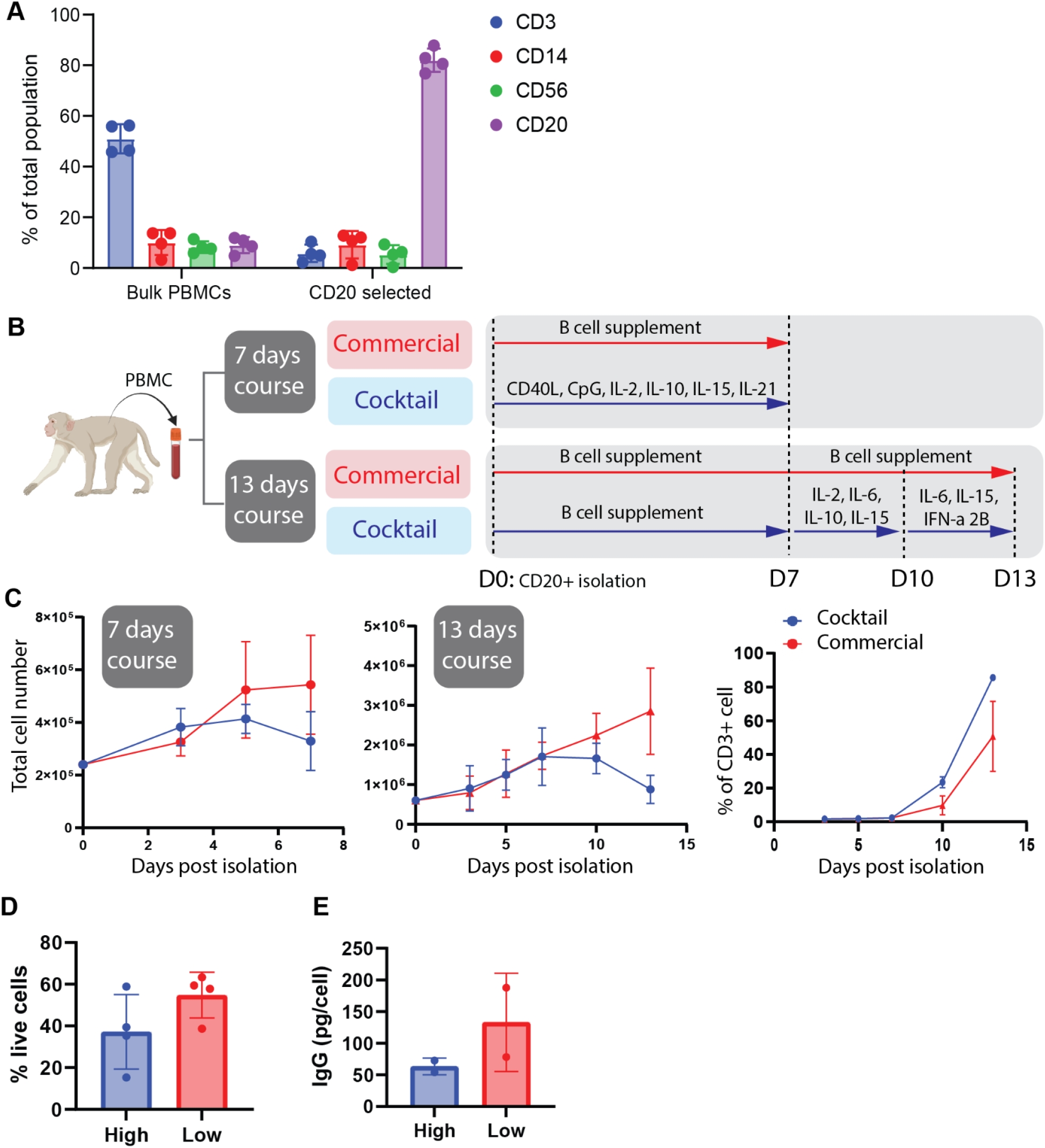
Method of ex vivo expansion and differentiated NHP PCs. **A.** Following isolation of CD20+ Rhesus B cells from PBMCs, we assessed cell purity via flow cytometry using the indicated antibodies (4 donors, n=4). **B.** Schematic workflow of monkey *ex vivo* differentiated PC generation with two different time course and two different conditions. **C.** CD20+ NHP B cells were cultured for 7 days/13 days with defined cocktail or commercial expansion medium (1.5-1×106 cells/mL) (3 donors, n=3), Total cells were counted on days 3, 5, and 7 (left and middle). Percentage of contaminating T cells cells (CD3+, left). **D-E.** To assess the impact of cell density, CD20+ NHP B cells were cultured in the commercial B cell medium for 7 days (Plated and maintained 1.5×10^6^ cells/mL versus plated at 2.5×10^5^ cells/mL and maintained at 1×10^5^ cells/mL; 4 donors, n=3). At day 7 following isolation, the cells were analyzed using flow cytometry. **D.** The percent viability, **E.** Supernatants and cells from the above experiment were assessed for IgG secretion via ELISA.

**Figure S3.**
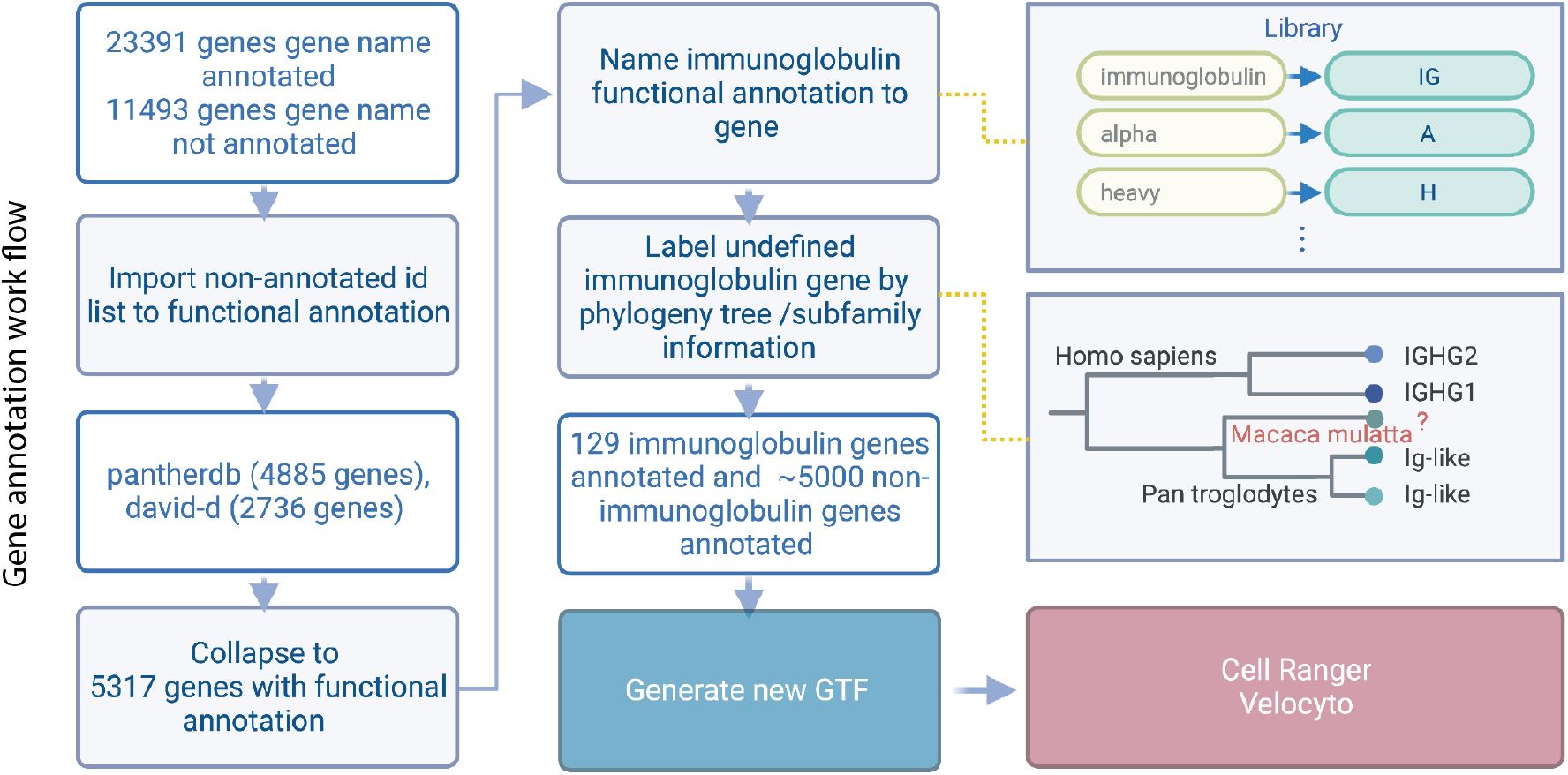
Monkey (Macaca mulatta) gene annotation workflow. Gene annotation workflow

**Figure S4:**
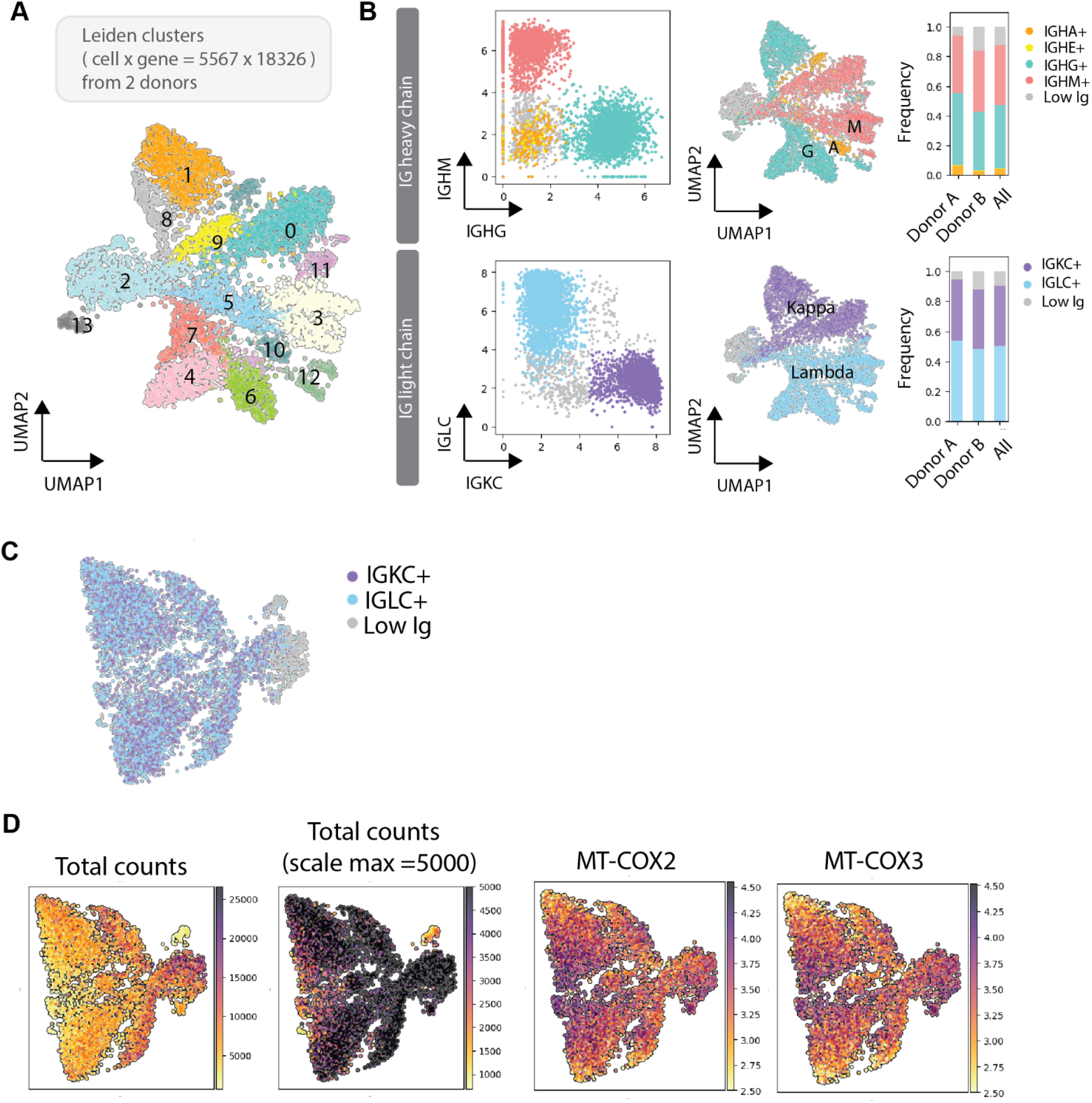
Monkey *ex vivo* differentiated PCs subsets classification and differential expression analysis for PCs. **A.** Single cell graph UMAP dimension-reduction projection of day13 B cells (n = 5567) from two biological replicates. **B.** Classification of immunoglobulin heavy chain (upper panel) and light chain (lower panel). UMAP projection of the subclass of IGHG+, IGHA+, IGHM+ cells (upper panel) and IGKC+, IGLC+ cells (lower panel), and bar graph represent the percentage of subclasses from two individual donor. **C.** UMAP projection of IGKC+, IGLC+ cells in LC-independent clusters. **D.** Heatmap showing total count and expression of representative MT genes in LC-independent clusters.

**Figure S5:**
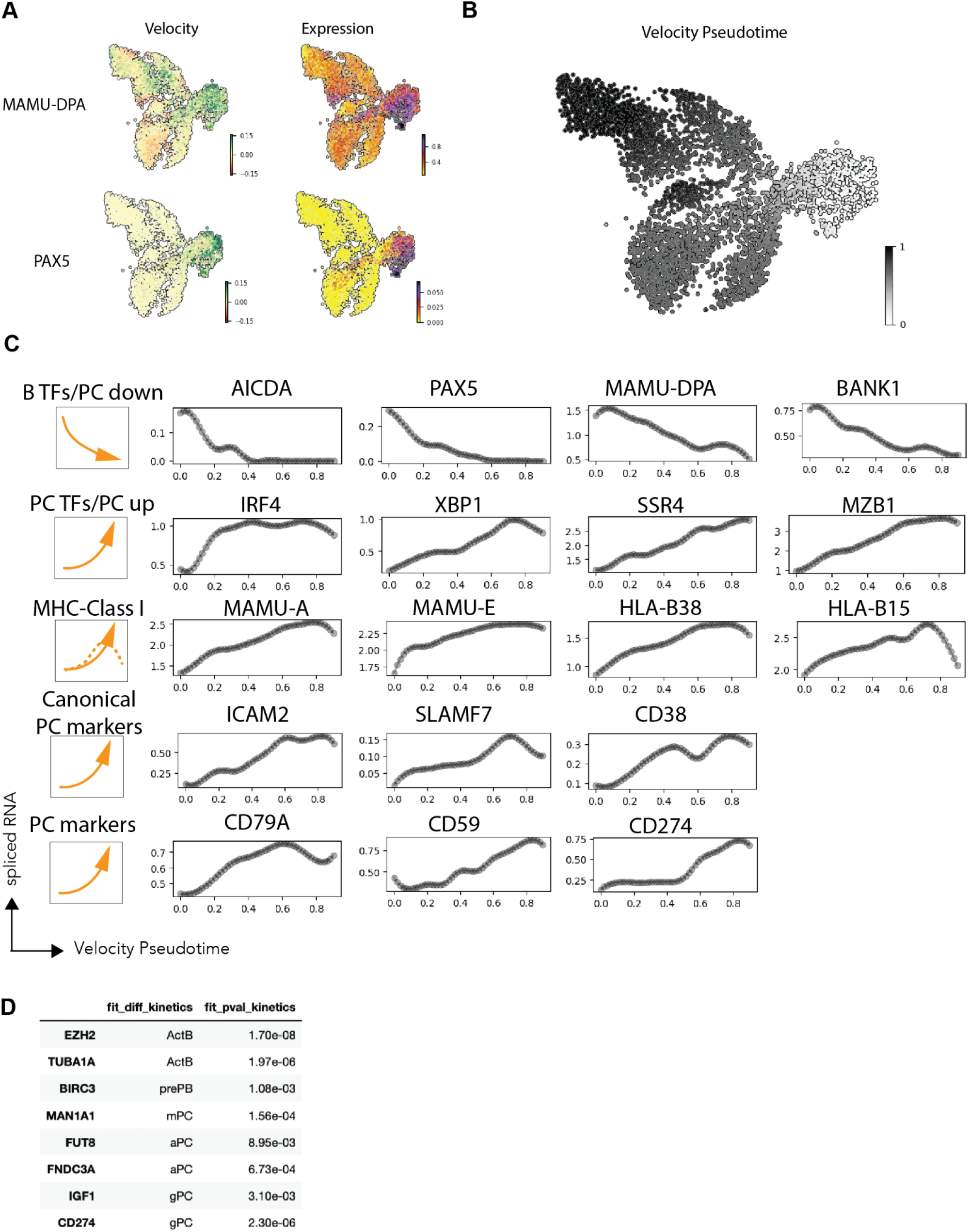
RNA velocity of immunoglobulin and kinetic differentiation analysis. **A.** Heatmap of indicated gene velocity and RNA expression level superimposed on to UMAP. **B.** Heatmap of velocity pseudotime superimposed on to UMAP. **C.** Scatter plot of indicated genes with velocity pseudotime and mean of spliced mRNA in every 0.1 pseudotime. **D.** scVelo kinetic differentiation analysis.

**Figure S6:**
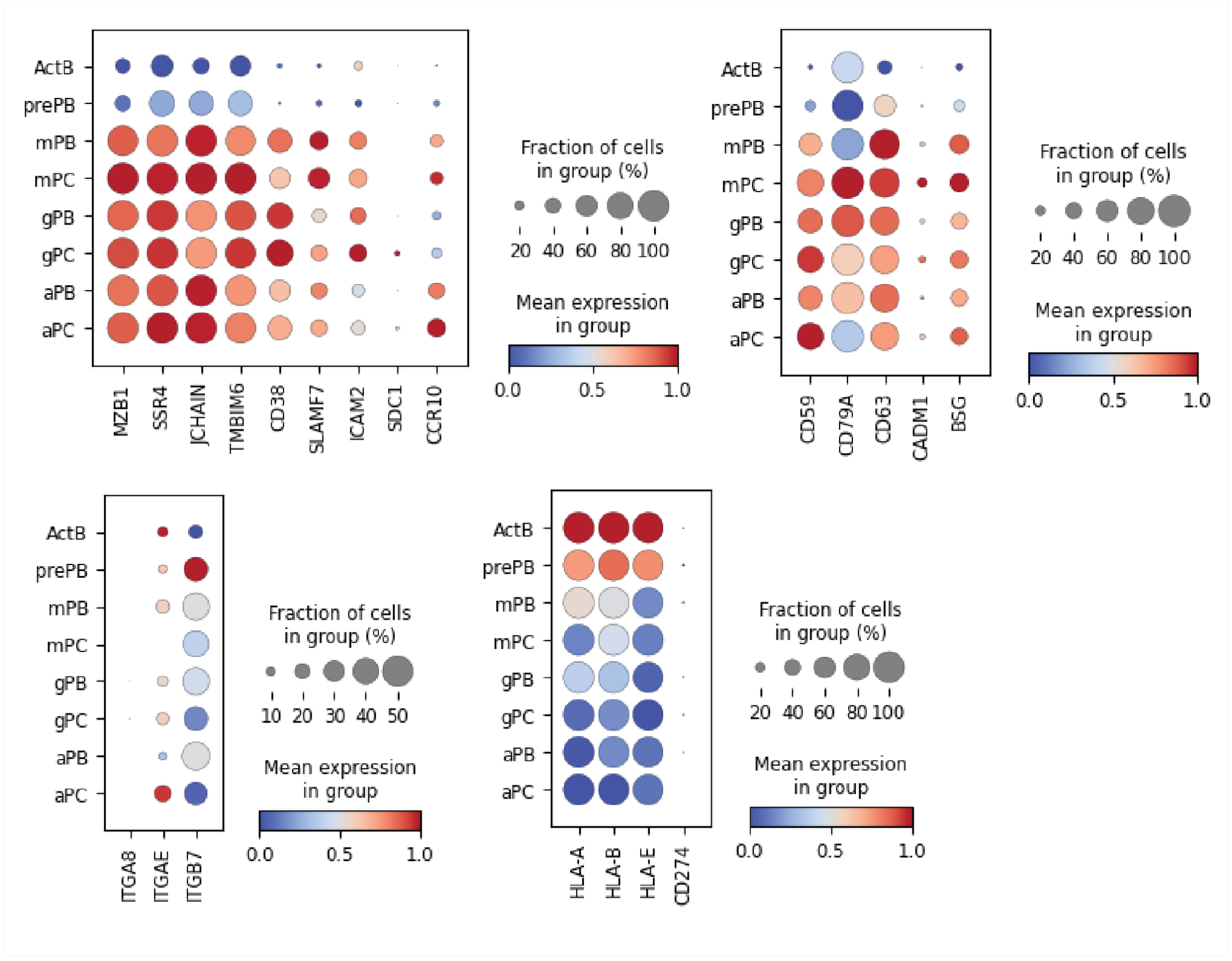
Comparison of human and monkey PC makers by scRNAseq analysis. Dotplot visualization of *ex vivo* differentiated human PC: subsets with different isotypes are listed on y-axis and representative genes from Fig. 3D are listed along the x-axis.

**Figure S7.**
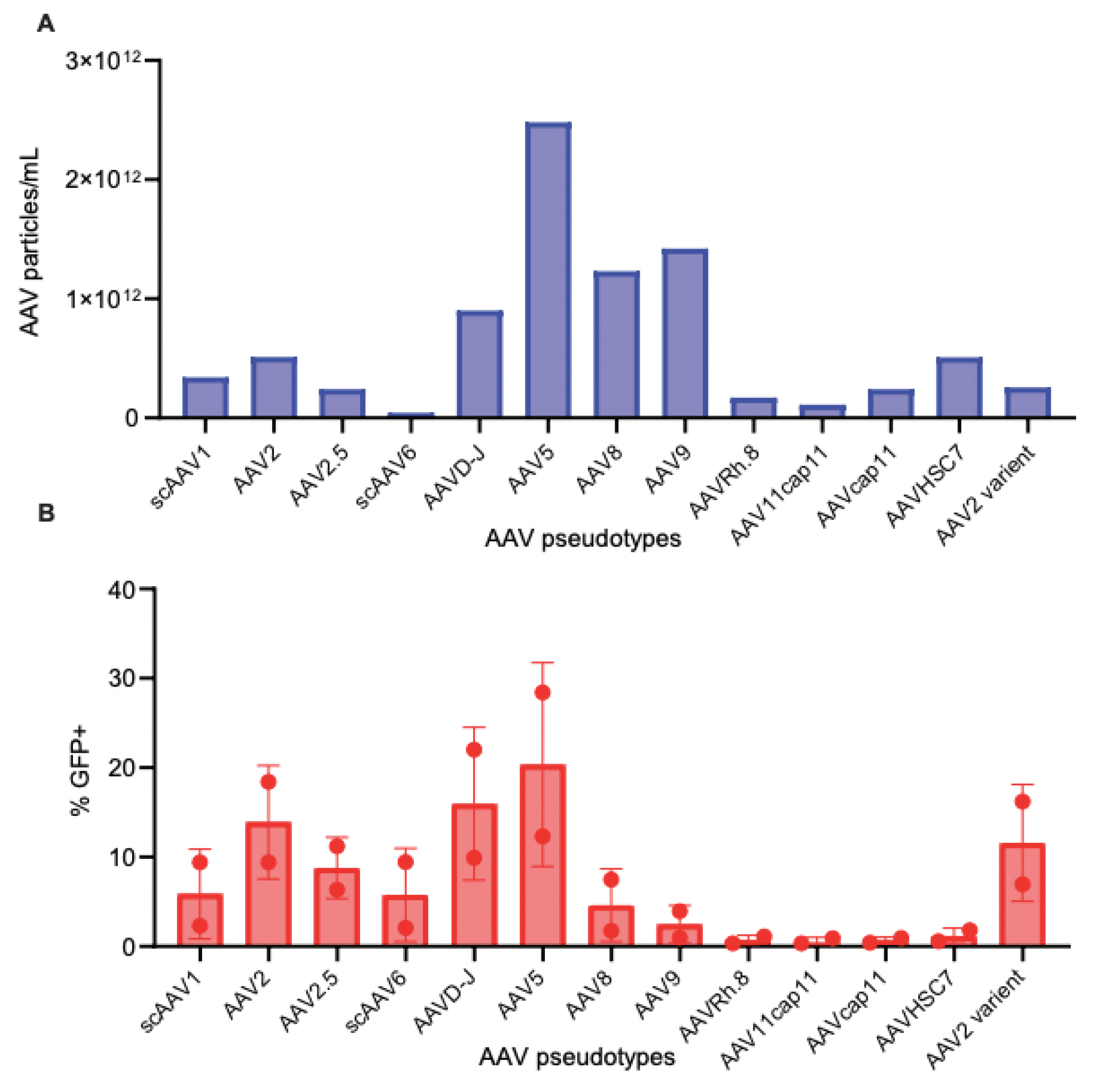
NHP B cell transduction with AAV. **A-B** NHP B cells were transduced on day 3 of culture. 20% by volume was added of GFP encapsulated in different AAV pseudotypes, with varying titers **A.** and flow for GFP+ **B.** was run 2 days later.

**Figure S8.**
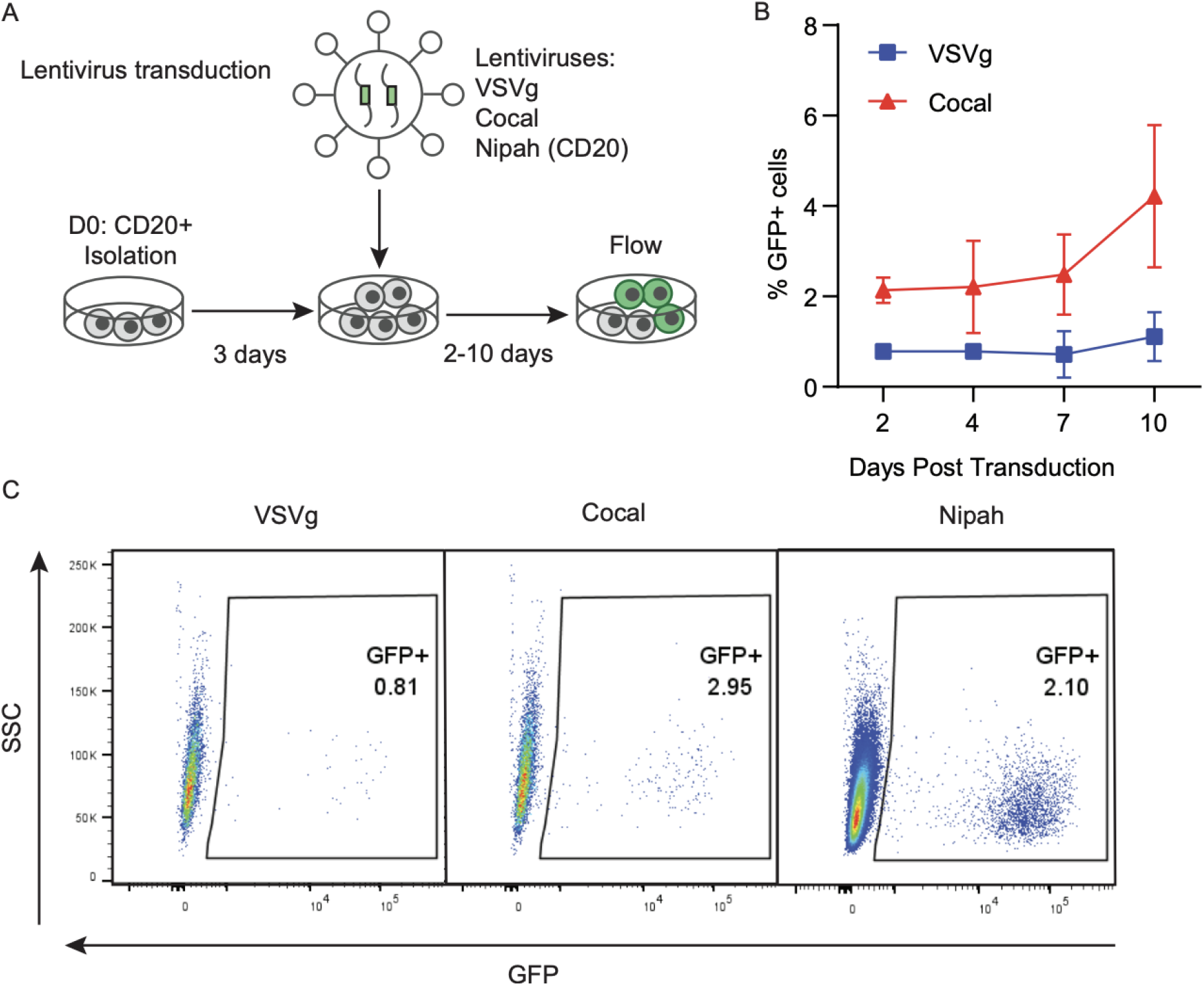
Lentiviral transduction of NHP B cells. **A.** Diagram of the lentiviral transduction and culturing protocol. **B-C** The percent GFP expression was quantified by flow cytometry in CD3-CD14-NHP B cells transduced with lentivirus pseudotyped with either VSVg or Cocal envelopes at an multiplicity of infection of 40 (2 donors, n=2) or with a Nipah anti-CD20 pseudotyped virus (1 donor, n=1). **B.** Graph showing GFP expression overtime for both VSVg and Cocal Pseudotypes **C.** Representative flow plots showing GFP percentage in CD3-CD14-cells 4 days post transduction with lentivirus pseudotyped with the indicated envelopes.

## Supplemental note: Nipah virus purification

To prepare the Nipah virus vector, 293T cells were plated in 10 CM plates per standard lentiviral preparation protocols. Each 10CM plate of 293T cells were transfected with transfer using polyethylenimine reagent as previously described (4 uL of reagent per 1 ug of DNA) and vectors using the following concentrations per plate: transfer vector (5 ug), PsPAX2 (5 ug), PMD2 Nipah F (1.5 ug) and PMD2 Nipah G (.3 ug). The Nipah F construct consists of amino acids 1-521 of the fusion protein from Henipavirus nipahense (GenBank sequence ID: NP_112026.1). The Nipah G construct consists of amino acids 35-602 of the attachment glycoprotein from Henipavirus nipahense (GenBank sequence ID: NP_112027.1) with the following changes (E468A, W471A, Q497A, and E500A), a linker sequence (SGGGGSGGGGSGGGGSGGSSAS), and amino acids 668-936 from the following vector sequence (GenBank sequence ID: ASW25860.1). One day following transfection, the media is changed (10% FBS in DMEM). Two days later, virus was collected and purified using ultracentrifiguation.

